# The hunt for ancient prions: Archaeal prion-like domains form amyloids and substitute for yeast prion domains

**DOI:** 10.1101/2020.07.20.212902

**Authors:** Tomasz Zajkowski, Michael D. Lee, Shamba S. Mondal, Amanda Carbajal, Robert Dec, Patrick D. Brennock, Radoslaw W. Piast, Jessica E. Snyder, Nicholas B. Bense, Wojciech Dzwolak, Daniel F. Jarosz, Lynn J. Rothschild

**Author notes:** M.D.L. and S.S.M. contributed equally to this work. **Classification** Biological Sciences: Evolution. **Author Contributions** T.Z., M.D.L., S.S.M., and L.J.R. designed research; T.Z., M.D.L, S.S.M, A.C., and R.D. performed research; T.Z., M.D.L., S.S.M., A.C., R.D., P.D.B., R.W.P., J.E.S., N.B.B., and W.D. analyzed data; T.Z., M.D.L., S.S.M., A.C., R.D., P.D.B., W.D, D.F.J., and L.J.R. wrote the paper.

## Abstract

Prions are proteins capable of acquiring an alternate conformation that can then induce additional copies to adopt this same alternate conformation. Although initially discovered in relation to mammalian disease, subsequent studies have revealed the presence of prions in Bacteria and Viruses, suggesting an ancient evolutionary origin. Here we explore whether prions exist in Archaea - the last domain of life left unexplored with regard to prions. After searching for potential prion-forming protein sequences computationally, we tested candidates *in vitro* and in organisms from the two other domains of life: *Escherichia coli* and *Saccharomyces cerevisiae*. Out of the 16 candidate prion-forming domains tested, 8 bound to amyloid-specific dye, and six acted as protein-based elements of information transfer, driving non-Mendelian patterns of inheritance. We additionally identified short peptides from archaeal prion candidates that can form amyloid fibrils independently. Candidates that tested positively in our assays had significantly higher tyrosine and phenylalanine content than candidates that tested negatively, suggesting that the presence of these amino acids may help distinguish functional prion domains from nonfunctional ones. Our data establish the presence of amyloid-forming prion-like domains in Archaea. Their discovery in all three domains of life further suggests the possibility that they were present at the time of the last universal common ancestor (LUCA).

**Significance Statement:** This work establishes that amyloid-forming, prion-like domains exist in Archaea and are capable of vertically transmitting their prion phenotype – allowing them to function as protein-based elements of inheritance. These observations, coupled with prior discoveries in Eukarya and Bacteria, suggest that prion-based self-assembly was likely present in life’s last universal common ancestor (LUCA), and therefore may be one of the most ancient epigenetic mechanisms.

## Introduction

One of the most notable and puzzling disease outbreaks of the last 50 years was bovine spongiform encephalopathy. Its baffling patterns of transmission arose because it was caused by a mysterious agent devoid of nucleic acids (1). The disease left 177 people and over 4 million cattle dead, harmed the economy of the UK, and, for a time, left the scientific community with no mechanism to explain the phenomenon (2). The cause was eventually determined to be a misfolded form of an endogenous protein, designated the Prion Protein (PrP) (1).

Although initially named for “proteinaceous infectious particles” (1), ascribing a stable, concise definition to the term “prion” is not trivial. As our scientific understanding of prion-like entities continues to develop alongside the concept’s integration into society’s lexicon, the term itself is under relatively rapid linguistic evolutionary pressure. One generalizable and useful definition is: *prions are proteins that convert between structurally and functionally distinct states, at least one of which is self-propagating and self-perpetuating* (slightly modified from Alberti *et al.*, 2009 (3) and Garcia and Jarosz, 2014 (4)). And although certainly useful, this does not adequately convey the whole story because it assigns the property of prion to the protein itself as an individual unit, while some of the key hallmarks of prion activity (e.g., self-propagation and non-Mendelian inheritance) only manifest through the interaction of multiple copies of a protein under specific conditions. Conceptually this is not entirely dissimilar from how quorum sensing can be thought of as an emergent property, rather than being ascribed to a single unit. It can therefore be useful to also think of the term “prion” as a state that emerges when a specific suite of characteristics is met for multiple copies of a protein that is capable of facilitating that state. That said, unless noted otherwise, herein the term “prion” will be utilized in the former, generalized form, referring to an individual protein.

While prions were being studied in mammals, puzzling non-chromosomal genetic elements were concurrently being discovered in yeast: [URE3] (5) and [*PSI*^+^] (6). As the prion story unfolded, it soon became clear that [URE3] and [*PSI*^+^] were prions of the Ure2p and Sup35p proteins, respectively (7–11). All known mammalian prion diseases are the result of the misfolding of the endogenous prion protein (PrP). By contrast, fungal prions are more diverse, sparking phenotypes produced by the alternative folds of many different proteins (7, 12–15). Prion conversion of PrP is pathogenic, whereas acquisition of many yeast prions is not. As more work has been done to elucidate nonpathogenic prion-forming proteins, it has been shown that prion conversion can serve as a molecular switch for multiple biological functions, often exerting a strong influence on survival. They can play a role in adaptive responses to environmental fluctuations (14, 16, 17), contribute to evolvability by acting as epigenetic elements (18–20), and act as evolutionary capacitors (21, 22) and bet-hedging devices (23–25). For example, *Saccharomyces cerevisiae* employs the [*GAR*^+^] prion to switch between specialist and generalist carbon-source utilization strategies, a switch that is heritable (26, 27). The [*ESI*^+^] prion drives the emergence and transgenerational inheritance of an activated chromatin state that can result in broad resistance to environmental stress, including antifungal drugs (28). The [*SMAUG*^+^] prion allows yeast to anticipate nutrient repletion after periods of starvation, providing a strong selective advantage (17). Other non-detrimental prions were found in many distinct species. In *Podospora anserina,* prion [Het-s] carries out normal cell functions in the process of self/non-self recognition (29). In the angiosperm *Arabidopsis thaliana*, the Luminidependens protein can form a prion and is responsible for signaling flowering as a result of detecting the temperature change in the environment (30). In baculoviruses, the prion forming LEF-10 protein was found to be responsible for efficient viral replication and expression (31). The discovery of functional prions across a broad phylogenetic diversity of organisms raises the question of their evolutionary origins.

The first prions to be discovered form highly ordered aggregates called amyloids (32) and many, but not all, currently known prions adopt an amyloid conformation. Amyloids are long, unbranched protein aggregates characterized by a fibrillar morphology and cross β-sheet quaternary protein structure (33). Just like prions, amyloids were first discovered in association with neurodegeneration. The most well-known amyloids form protein deposits in the human brain as in the cases of Aβ peptide and Tau protein in Alzheimer’s disease, huntingtin in Huntington’s disease, α-synuclein in Parkinson’s disease, and SOD1 in Amyotrophic lateral sclerosis (34). Similar to prions, amyloids exist in myriad organisms, including microbes. Although their presence correlates with neurodegeneration, some have argued that amyloid itself might be overall more beneficial relative to other oligomeric forms of aggregation-prone proteins (35, 36). Amyloids have functional and nonpathogenic roles in species as phylogenetically distinct as *Escherichia coli* and humans (37, 38). The term “functional amyloids” was created to describe non-detrimental amyloids (39). In microbes, functional amyloids are often found in the extracellular matrix contributing to cellular protection and biofilm formation (38, 40). The first functional amyloid identified in humans was the PMEL 17 protein (39, 41). Important in melanogenesis, amyloids of PMEL 17 create a scaffold for melanin distribution (39, 41). Amyloid-forming proteins have been detected and experimentally verified in all domains of life: Eukaryota, Bacteria, and Archaea (42–44). Prions, on the other hand, have to date only been discovered and verified in Eukarya and Bacteria (1, 45).

The propagation of known prions relies on a protein fragment called the prion domain (PrD). These domains are often enriched in glutamine and asparagine residues and are usually found in the disordered fragments of proteins. Leveraging these principles, bioinformatic algorithms can be used to predict novel PrDs (46–48). Here we use computational prediction methods on 1,262 archaeal proteomes to assess the distribution of prion proteins across the archaeal domain. We identify candidate PrDs (cPrDs) and investigate the functions associated with the proteins harboring them. We additionally perform a suite of biochemical and genetic experiments on a subset of our identified cPrDs to validate their capacity to form infectious aggregates. Our data reveal that multiple archaeal proteins can act as prions, forming heritable protein aggregates that are capable of acting as protein-based epigenetic elements. These observations, coupled with prior discoveries in eukarya and bacteria, suggest that prion-based self-assembly is an ancient form of epigenetics that may have been present in life’s last universal common ancestor (LUCA).

## Results

### Computational identification and distribution of cPrDs in Archaea

Leveraging the strong enrichment of glutamine and asparagine in prion domains, we utilized the program PLAAC (49) to scan for cPrDs in archaeal proteomes sourced from the UniProt (50) database (see Fig. 1 for overview). Of the 1,262 archaeal proteomes scanned, at least one cPrD was identified in 873 proteomes (~69%). Of the 2,805,234 proteins scanned, 2,797 were identified as containing cPrDs (~0.10%). The distribution of proteomes with at least 1 cPrD identified was largely homogenous across the archaeal domain (Fig. 2). There were modestly lower proportions within the Bathyarchaeota (58%) and the Asgard group (62%), and a higher proportion within the Crenarchaeota (79%). We note however that these taxonomic summary delineations are somewhat arbitrary, and as with all databases there can be underlying confounding factors about which lineages may be over- or under-represented as compared to others. For example, a commonly sequenced organism will likely be overrepresented, as compared to those less commonly sequenced.

**Fig. 1.**
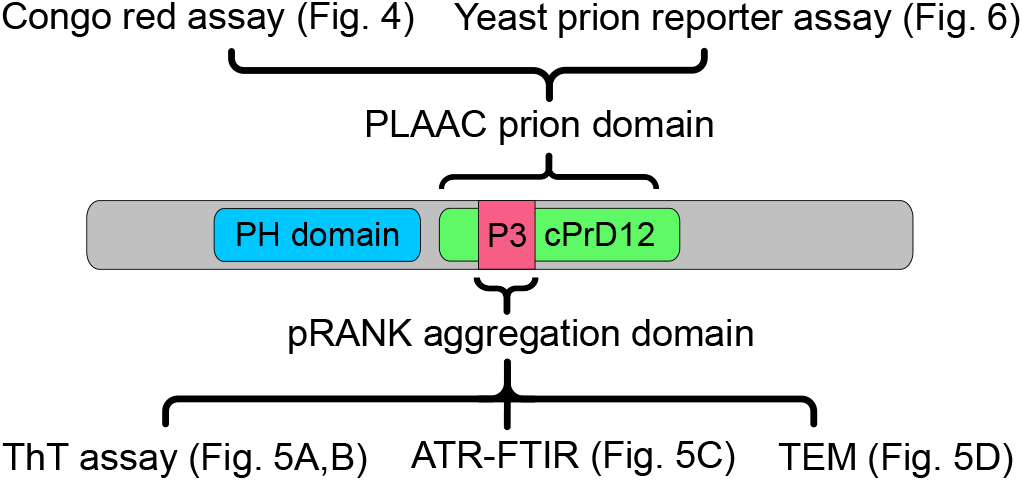
Fragments of protein and methods used to analyze them. Schematic representation of PH domain-containing protein from *Sulfolobus* sp.. Candidate prion domain identified by PLAAC (cPrD12) and aggregation domain identified by pRANK (P3) are marked. Methods applied to analyze the designated fragments are listed with references to other figures.

**Fig. 2.**
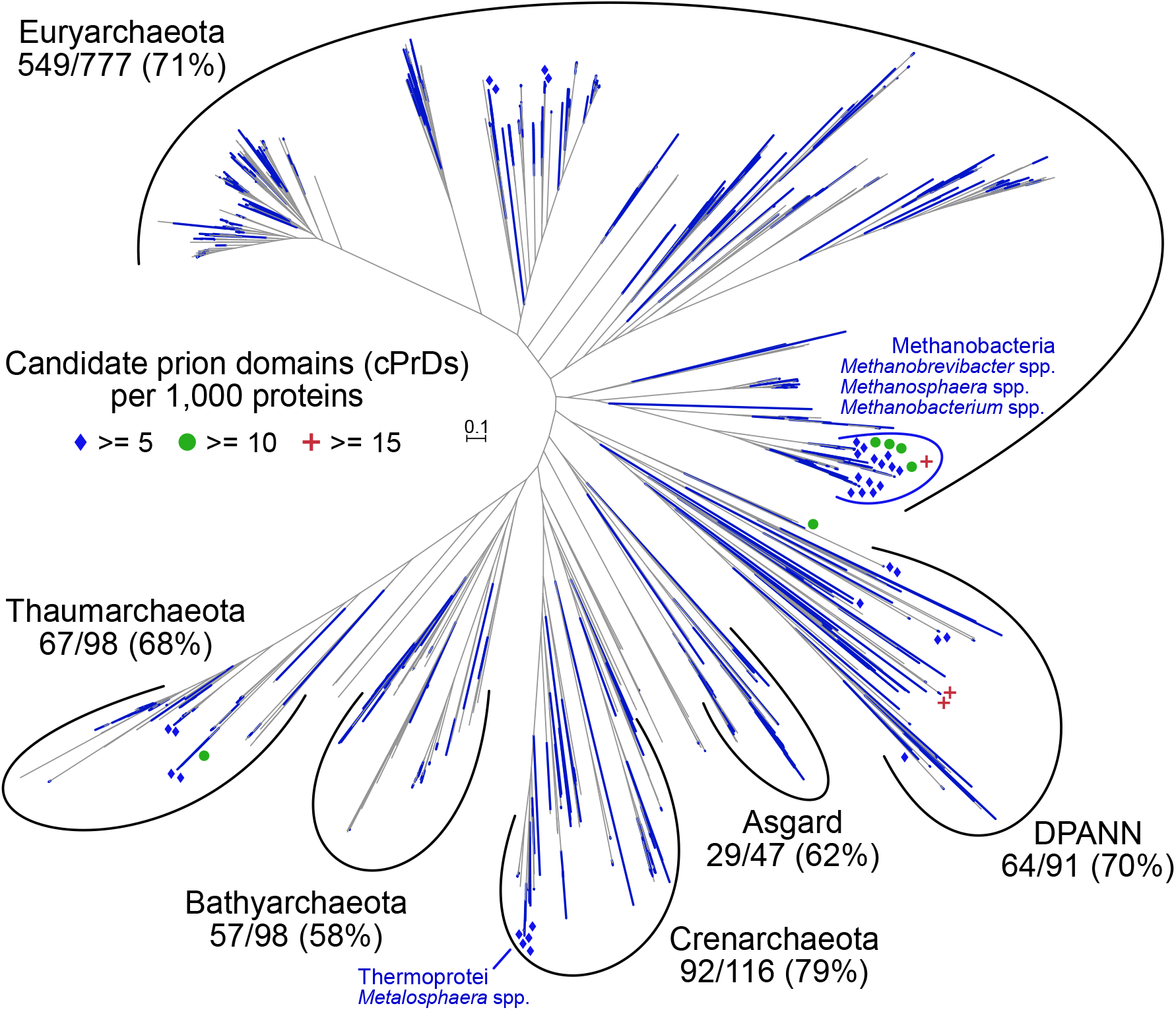
A Phylogenomic tree of incorporated Archaea with distribution of cPrDs overlain. Tips with at least 1 cPrD are colored blue. Blue diamonds are places near those with ≥ to 5 cPrDs per 1,000 genes; green circles near those with ≥ 10; and red pluses near those with ≥ 15. Major taxa summary values include the number within that group with identified cPrDs out of the total in that group followed by that percentage. The Asgard group here includes: Thorarchaeota, Odinarchaeota, and Heimdallarchaeota. DPANN group here includes: Woesearchaeota, Pacearchaeota, Nanoarchaeota, Aenigmarchaeota, Diapherotrites, Micrarchaeota, and Altiarchaeota. The unrooted tree is an estimated maximum-likelihood of 76 single-copy orthologs forming an alignment of 12,330 amino acids (see Methods).

The frequency of identified cPrDs within each proteome was ~2.21 ± 3.38 (mean ± 1SD) with a median of one: 30% of the scanned proteomes contained 0; ~25% had the median of 1: ~35% had between 2-5; and ~10% had greater than 5 (Supp. Dataset 1). Normalizing these values to the total number of proteins in each individual proteome (done as cPrDs per 1,000 proteins) returns ~1.14 ± 1.89 (mean ± 1SD) with a median of 0.6 (Supp. Dataset 1; histogram in Supp. Fig. S1). Based on these normalized values, 3 proteomes were found to have > 15 identified cPrDs per 1,000 genes (placing them in percentile ranks of > 99.8) including 2 closely related Nanoarchaeota (GCA_003086475.1 and GCA_003086415.1) and a Euryarchaeota of the class Methanobacteria (GCA_001639265.1; demarcated with red plus signs in Fig. 2). Other proteomes with relatively high-densities of identified cPrDs included 10 with ≥10 cPrDs per 1,000 proteins (99.2 percentile rank) and 46 with ≥ 5 (96.4th percentile). Many of these were concentrated within the class Methanobacteria (across the genera *Methanobrevibacter*, *Methanosphaera*, and *Methanobacterium*) and within representatives of *Metallosphaera sedula* of the Thermoprotei class (Fig. 2).

### Enriched functions of proteins containing cPrDs

We utilized Gene Ontology (GO) (51) annotations (GO “terms”) and goatools (52) to test for enrichment or purification of specific GO terms in our identified cPrD-containing proteins (2,797) as compared to all of the proteins scanned (2,805,234). The identified cPrD-containing proteins were associated with 706 GO terms, 523 of which were found to be significantly enriched or purified in the cPrD group relative to all proteins (based on Benjamini-Hochberg false-discovery rates ≤ 0.05; Supp. Dataset 2). The vast majority of these (469/523; ~90%) were found to be purified in proteins harboring cPrDs relative to all proteins (under-represented), and 54 were found to be enriched – a subset of which are presented in Fig. 3.

**Fig. 3.**
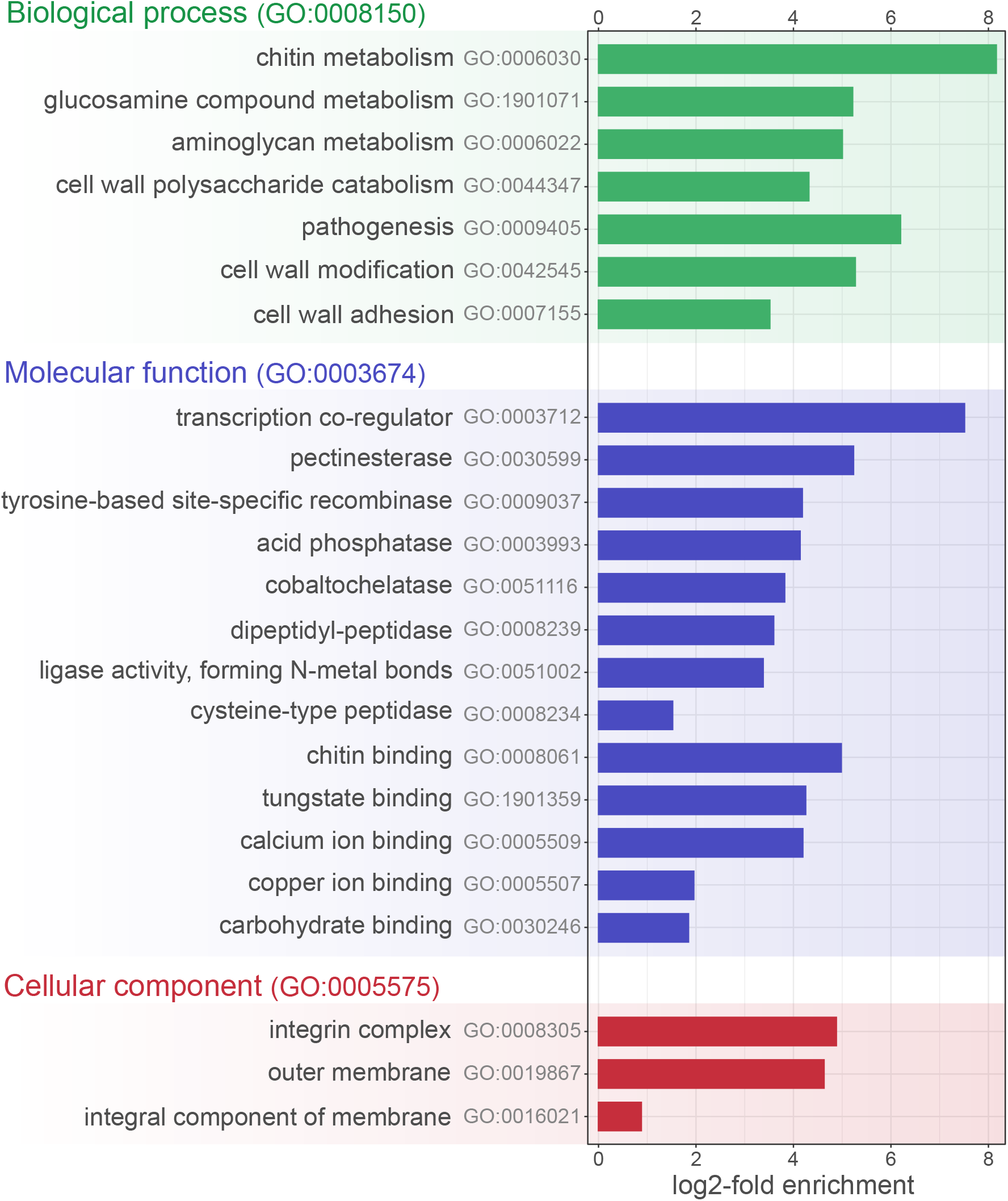
Select enriched GO terms of proteins harboring cPrDs as compared to all scanned archaeal proteins. Displayed are log2-fold enrichments of the given GO term’s frequency among total cPrD-containing proteins (2,797) as compared its frequency among all computationally scanned archaeal proteins (2,805,234). Considered enrichments were those with Benjamini-Hochberg false-discovery rates of ≥ 0.05 (see Methods). Full results in Supp. Dataset 3.

GO breaks its terms down into 3 namespaces: Biological Process, Molecular Function, and Cellular Component. Significantly enriched GO terms included proteins involved in: chitin metabolism (GO:0006030), cell wall polysaccharide catabolism (GO:0044347), and cell wall adhesion (GO:0007155) within the Biological Process namespace; transcription co-regulator (GO:0003712), tyrosine-based site-specific recombinase (GO:0009037), and calcium (GO:0005509) and copper ion binding (GO:0005507) within the Molecular Function namespace; and outer membrane (GO:0019867) within the Cellular Component namespace (Fig. 3; full results in Supp. Dataset 2). Significantly purified (under-represented) GO terms included: cellular amino acid (GO:0008652) and nucleotide biosynthesis (GO:0009165) within Biological Process; nucleotide (GO:0000166) and metal ion binding (GO:0046872) within Molecular Function; and cytoplasm (GO:0005737) and ribosome (GO:0005840) within Cellular Component (Supp. Dataset 2).

### Multiple archaeal cPrDs bind Congo red

Selective binding to dyes, including Congo red, is a diagnostic characteristic of amyloid. We investigated this property in the cPrDs of the highest scoring (COREscore ≥30) putative prions from archaeal proteomes. Many cPrDs consisted of sequences with long stretches of only Q or N. Because of the low complexity of these sequences, the synthesis of coding DNA in many cases proved impossible. Thus, we chose sequences complex enough to allow their synthesis. From this group, we rejected genes with unknown functions, and synthesized the remaining 16 coding sequences (Table 1). We cloned these sequences into a commercially available vector allowing synthesis and export of investigated cPrDs (53).

**Table 1.**

| ID | Congo red assay, results | Yeast prion reporter assay, results | NCBI Accession | NCBI Taxonomy | NCBI Annotation |
| --- | --- | --- | --- | --- | --- |
| cPrD1 | CR+ | YPR+ | AEH61342.1 | *Methanosalsum zhilinae* | transcriptional repressor, CopY family |
| cPrD2 | CR- | YPR- | AGN16545.1 | *Methanobrevibacter* sp. | adhesin-like protein |
| cPrD3 | CR- | YPR+ | ARM76690.1 | *Acidianus manzaensis* | zinc ribbon domain-containing protein |
| cPrD4 | CR+ | YPR- | KZX15367.1 | *Methanobrevibacter filiformis* | tRNA biosynthesis protein MnmC |
| cPrD5 | CR+ | YPR- | OQD58440.1 | *Methanobrevibacter arboriphilus* | putative peptidase |
| cPrD6 | CR- | YPR+ | WP_009487818.1 | *Halobacterium* sp. | DNA starvation/stationary phase protection Dps |
| cPrD7 | CR- | YPR+ | WP_013774943.1 | *Acidianus hospitalis* | zinc ribbon domain-containing protein |
| cPrD8 | CR+ | YPR+ | WP_014027445.1 | *Pyrolobus fumarii* | Tyrosine-type recombinase/integrase |
| cPrD9 | CR+ | YPR+ | WP_042706453.1 | *Methanomicrobium mobile* | YIP1 family protein |
| cPrD10 | CR- | YPR+ | WP_054838334.1 | *Sulfolobus metallicus* | zinc ribbon domain-containing protein |
| cPrD11 | CR+ | YPR+ | WP_066970924.1 | *Methanobrevibacter filiformis* | NADAR family protein |
| cPrD12 | CR+ | YPR+ | WP_066972153.1 | *Methanobrevibacter filiformis* | PH domain-containing protein |
| cPrD13 | CR+ | YPR+ | WP_069282784.1 | *Sulfolobus* sp. | zinc ribbon domain-containing protein |
| cPrD14 | CR- | YPR- | WP_074791546.1 | *Haloferax larsenii* | ribonuclease BN |
| cPrD15 | CR- | YPR- | WP_082223394.1 | *Halosimplex carsbadense* | CBS domain-containing protein |
| cPrD16 | CR- | YPR- | WP_084269446.1 | *Methanobrevibacter curvatus* | ATP-dependent protease LonB |

To test if the putative prions were capable of forming amyloid fibrils, we used a bacterial export system for generating extracellular amyloid aggregates (53). The use of this system enables rapid screening for the ability of the aggregates to bind to the Congo red dye. A full-length prion protein is not always necessary, nor optimal for prion formation (54, 55). For example, in most yeast prions proteins, overproduction of only the amyloid-forming portion of the protein is sufficient for prion induction (8, 56). This induction is sometimes more efficient than with overproduction of the full-length protein (57, 58). Another characteristic of known PrDs is their modularity, which allows them to be cloned into other proteins thus transferring the ability to form prion aggregates (47). Preliminary tests also demonstrated that the use of a cPrD increases the sensitivity of the Congo red assay compared to full-length proteins (Supp. Fig. S2). For these reasons, and synthesis considerations, we tested cPrDs instead of full-length proteins.

Bacteria transformed with plasmids encoding the cPrDs were plated on an induction medium containing Congo red. Because in the C-DAG system the expressed cPrDs are exported outside of the cell they can interact with the dye in the culture medium. Bacterial colonies that produce cPrDs capable of forming amyloids bind Congo red, which then changes the color of the colony to red. A clear change in color was observed in 8 out of 16 colonies (Fig. 4; see Supp. Fig. S3 for results of all 16).

**Fig. 4.**
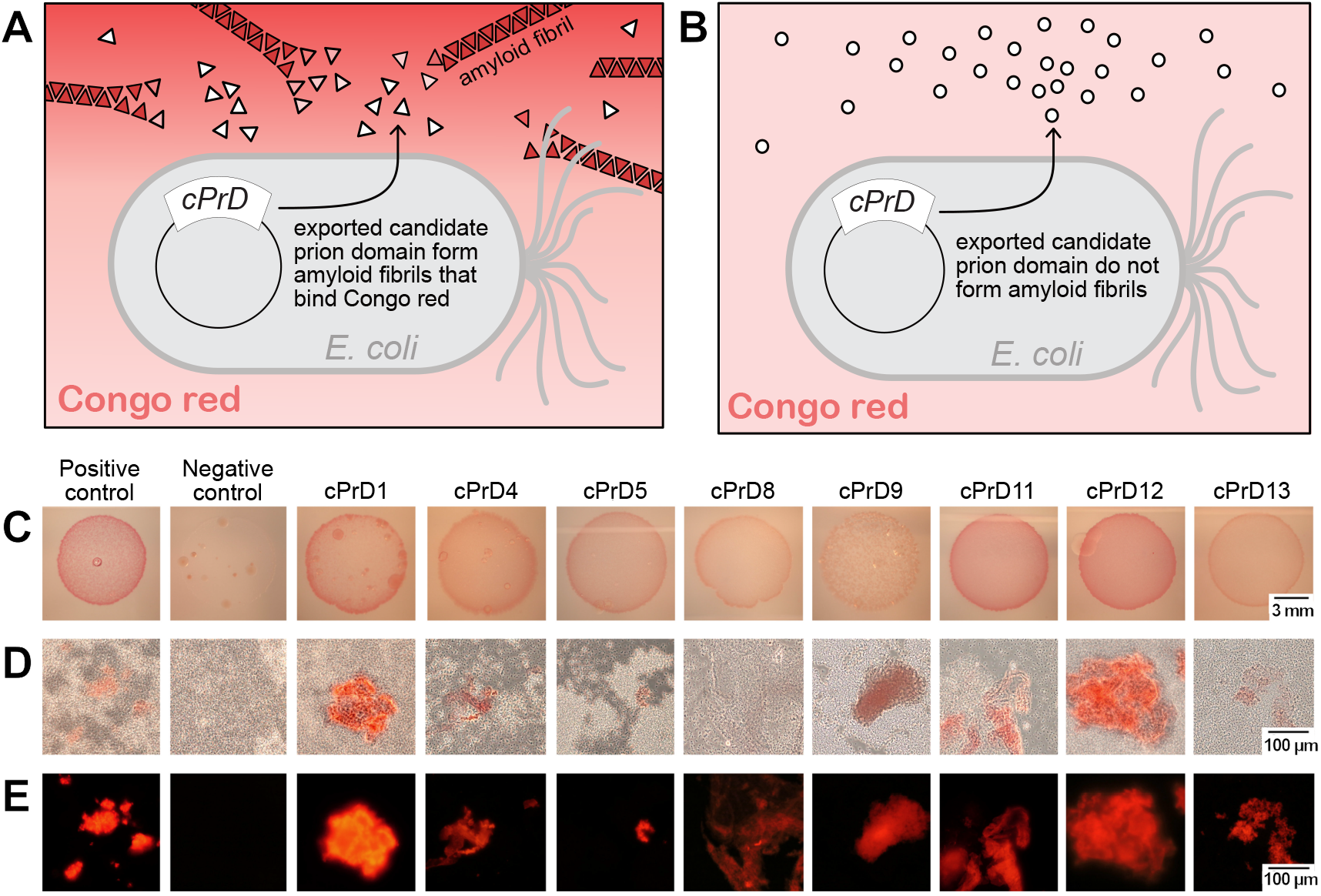
Comparison of colonies and corresponding protein aggregates produced by bacteria transformed with plasmids encoding cPrDs. A. and B. Two variants of possible results described in a cartoon. **A.** Candidate cPrD forms amyloid aggregates that bind Congo red. **B.** cPrD does not form amyloid aggregates therefore does not bind Congo red. **C.** Colonies of *E. coli* grew on the induction medium containing 0.1% Congo red. The red color of the colony indicates that the exported protein binds to Congo red. Binding of Congo red is typical for amyloid fibrils. **D.** Light microscopy of preparations made of the same bacterial colonies show aggregates binding Congo red and bacterial cells concentrating around them. **E.** Fluorescent microscopy of protein aggregates from panel B is shown. The amyloidogenic fragment of Sup35 (residues 2-253) was used as a positive control. Fragment of Sup35 not able to form amyloids (residues 125-253) was used as a negative control. Protein names and organisms from which candidate prion domains (cPrDs) were selected are listed in table 1.

Microscopy samples were prepared from the colonies that obtained the strongest color (Fig. 4D). Using light microscopy, we confirmed the presence of Congo red binding aggregates in 8 samples as well as in positive controls, but saw no binding to negative controls. We observed that *E. coli* cells tend to concentrate around the aggregates, consistent with the known role of amyloid aggregates in cell adhesion in biofilms (59). Detection of Congo red fluorescence in some cases made aggregates more noticeable (Fig. 4E). In summary, we showed that several cPrDs from Archaea can bind amyloid specific dye. This is the first step in verification of possible prion nature of the proteins.

### Archaeal cPrDs form amyloids *in vitro*

Congo red staining alone is insufficient for confirmation of the amyloid nature of protein aggregates (60). Thus, to further evaluate the ability of archaeal prion candidates to form amyloids, we employed an additional set of tests. Following the hypothesis that small regions of a protein are often both necessary and sufficient to drive prion behavior (58), we analyzed even further shortened segments of the cPrDs. The length of cPrDs that yielded positive results in the Congo red assay varied from 62 to 179 amino acids. To choose fragments of putative archaeal prions most likely to drive aggregation, we used pRANK that assigns 20 amino acid long sequence window. As a principle, we selected for further analysis only these peptides which were initially soluble in water. We observed a very significant increase in thioflavin T (ThT) fluorescence for aggregated forms of peptides P1, P4 and moderate increase of ThT emission for P2 and P3 (Fig. 5). Binding of ThT to amyloids increases the quantum yield of fluorescence of the dye (61). Varying enhancement of ThT fluorescence is often observed for amyloid fibrils, depending on structural features that may reduce ThT binding affinity (62). Furthermore, we found that dissolving some peptides in water will trigger a spontaneous increase in ThT emission intensity until a plateau is reached (e.g. P4 - see Fig. 5B), a hallmark of amyloid formation. To examine the secondary structure of the aggregates, we used attenuated total reflectance Fourier transform infrared spectroscopy (ATR−FTIR) (Fig. 5C). The major spectral components of the vibrational amide I band were below 1630 cm^−1^ for all four peptides we tested, implying the presence of parallel beta-sheet structure characteristic of amyloid fibrils (63). For P1, the beta-sheet signal at *ca.* 1629 cm^−1^ flanks the dominant spectral component at 1656 cm^−1^ indicating possible coexistence of tightly packed amyloidal beta-core with abundant and less ordered conformations. A similar observation has been made for amyloid fibrils of Tau protein (64). We also performed a molecular dynamics simulation of peptide P2 showing that presence of a preformed β-sheet promotes the formation of a new filament of β-sheet out of unfolded P2 peptide (Supp. Fig. S4). Finally, to further examine the morphology of the aggregates, we performed transmission electron microscopy. These analyses confirmed the presence of amyloid fibrils in all analyzed preparations (Fig. 5D). Collectively these data establish that amyloid conversion is promoted by specific sequences within the cPrDs.

**Fig. 5.**
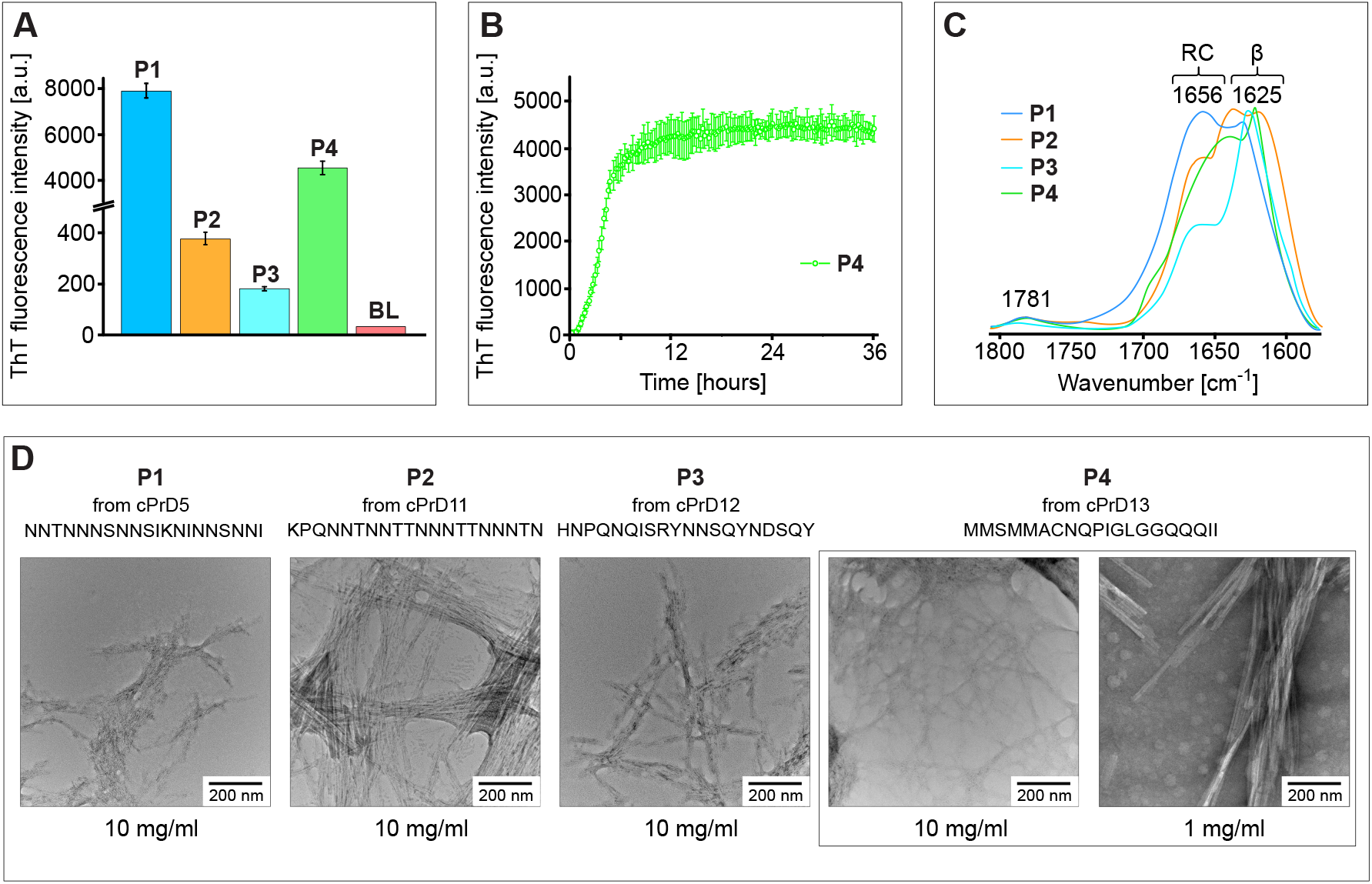
Analysis of aggregates formed by pRANK-peptides. **A.** Histogram showing ThT fluorescence intensity levels of samples containing peptides derived from prion candidates OQD58440.1 (P1); WP_066970924.1 (P2); WP_066972153.1 (P3); WP_069282784.1(P4). Prior to measurement, samples with the composition: 2.5 mg / mL peptide, 0.05 M NaCl, 20 μM Th, H2O, pH around 3, were incubated for 24 hours at 37 degrees. Three independent measurements were made for each sample, which was the basis for calculating the mean and error bar. For comparison, the result for the blank is also presented (BL). **B.** ThT fluorescence-monitored reassociation of WP_069282784.1 (P4) peptide. Sample composition and measurement conditions are the same as in the case of A. **C.** ATR−FTIR spectra of the amide I band region of samples containing pRANK-peptides. Before measurement, small portions of lyophilized peptides were suspended in water, which was then gently evaporated. **D.** Micrographs of all four peptides tested obtained using a transmission electron microscope show the fibrillar morphology of the aggregates. Peptide WP_069282784.1 (P4) is shown in two different concentrations.

### Archaeal cPrDs can promote protein-based inheritance

The unusual folding landscapes of prion proteins allow them to act as epigenetic mechanisms of inheritance (46). To test whether this was true for our identified archaeal cPrDs, we employed an assay based on the well-characterized prion phenotypes of *Saccharomyces cerevisiae* that depend on the prion state of Sup35 protein (3). Normally, the protein Sup35 is a part of a translation-termination complex responsible for recognition of the stop codon and termination of translation. The yeast strain that we used in these experiments has a premature stop codon in *ADE1* gene coding for SAICAR synthetase. Premature translation termination in *ADE1* causes accumulation of red pigment – the product of polymerization of aminoimidazole ribotide (AIR) (65, 66) – and the ensuing lack of adenine production precludes the growth of naive, [*psi*^+^] cells on adenine-deficient medium (SD-Ade). Occasionally, Sup35 undergoes conformational conversion to a prion state in which it forms amyloid fibrils. In this state, the protein is sequestered into aggregates, inhibiting translation termination - the [*PSI*^+^] phenotype - and promoting readthrough of the premature stop codon in *ADE1*. Colonies with this phenotype can grow on SD-Ade medium, and appear white on nutrient-rich media like Yeast Peptone Dextrose (YPD) because accumulation of the red pigment is reduced.

Sup35 protein consists of an N-terminal PrD, a middle region (M), and a C-terminal functional domain (C). The PrD of Sup35 and other prions are modular and can be transferred to non-prion proteins creating new protein-based elements of inheritance (47). In our experiment, we tested if the original PrD of Sup35 can be substituted by cPrDs from archaea while preserving the ability to undergo prion-dependent phenotypic change. If a cPrD can function as a *bona fide* prion domain, the yeast colony bearing the chimeric protein should have a phenotype resembling [*PSI*^+^]. In contrast, a cPrD that is unable to facilitate the prion-like conversion would have a [*psi*^−^] phenotype.

We substituted the original PrD of Sup35 with 16 different archaeal cPrDs, generating a series of cPrD-Sup35MC chimeric proteins. In this yeast-based prion reporter assay (YPR) we found that 10 chimeric strains grew colonies of a significant size on SD-Ade media. To distinguish quantitatively between positive (YPR+) and negative (YPR-) results of the assay, images of the Petri dishes were analyzed using ImageJ (see Methods for details). Most chimeric strains when plated on YPD medium grew colonies much whiter than the negative control, suggesting aberrant synthesis of red pigment (Fig 6; see Supp. Fig. S5, S7, S8.1, and S8.2 for all positive and negative results; results are also summarized in Table 1 and Supp. Dataset 3). The reduction of red pigment is characteristic for [*PSI*^+^] phenotype and indicates the presence of prion form of Sup35. We re-streaked the colonies that generated a phenotype resembling [*PSI*^+^] (YPR+) on YPD media. Some colonies showed reversion to [*psi*^−^] phenotype suggesting that cPrD-Sup35MC is in a metastable state allowing reversion of prion phenotype (Supp. Fig. S6).

**Fig. 6.**
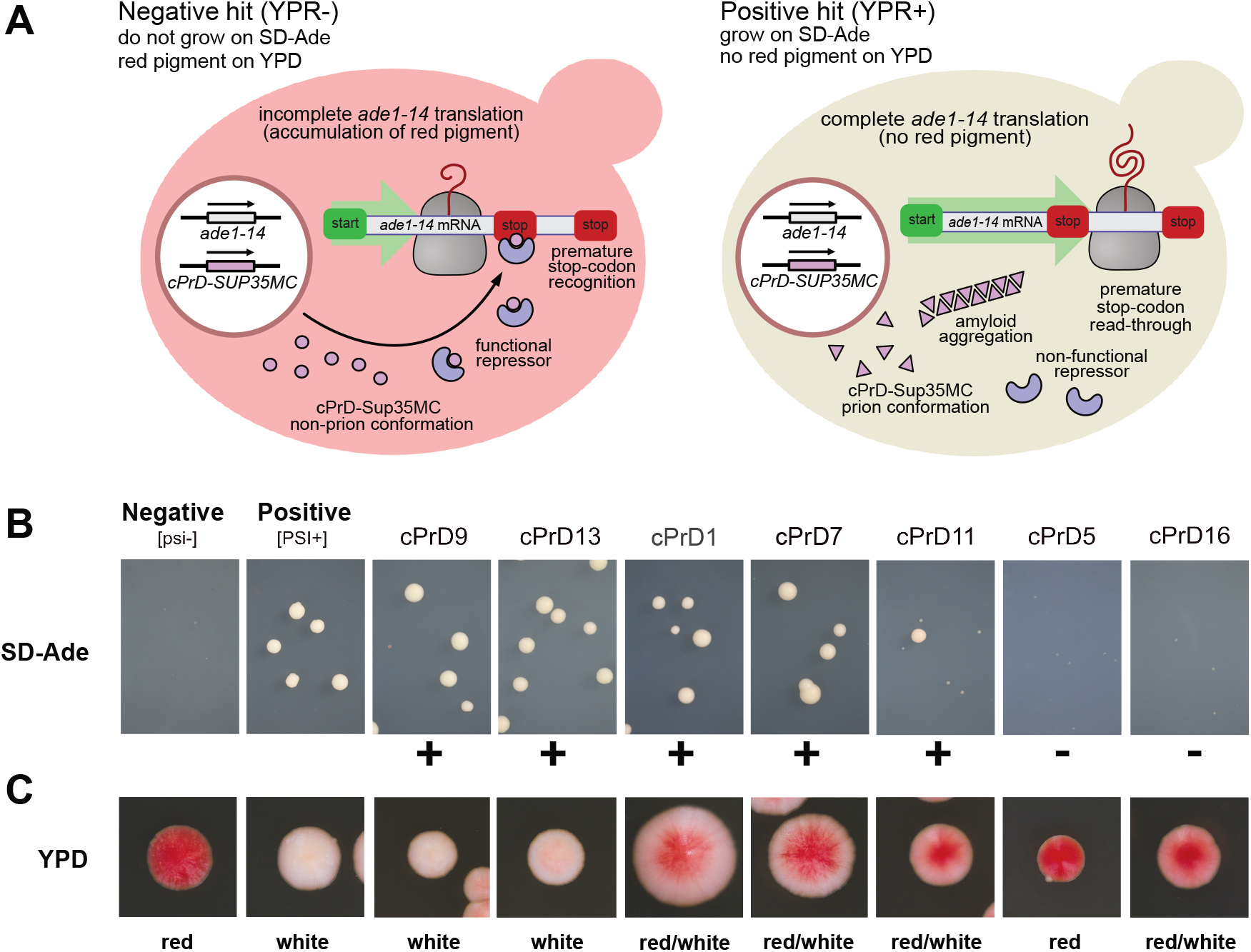
cPrD-Sup35MC strains display different ability to grow on media lacking adenine and a variety of colony colors. **A.** Cartoon explaining the mechanism of accumulation of red pigment, which depends on the formation of functional repressor. **B and C.** Representative images of the colonies of cPrD-Sup35MC-expressing strains growing on SD-Ade and YPD plates respectively (see Supp. Fig. 5 for all 16 tested and Supp. Fig. 7 for images of the whole area of Petri dishes). Colonies phenotypes [*psi*^−^] and [*PSI*^+^] are shown for comparison. Positive results are marked with “+” and are considered positive results of the yeast prion reporter assay (YPR+). Negative results are marked with “−” and are considered negative results of the yeast prion.

Based on this set of experiments we conclude that archaeal cPrDs can functionally substitute PrD of Sup35 and facilitate the formation of prion-based elements of inheritance.

### Tyrosine and/or phenylalanine content distinguishes functional cPrDs

In the experiments described above we found a total of six cPrDs that tested positive both in Congo red and yeast prion reporter assays (CR+/YPR+), and four that tested negative in both assays (CR−/YPR−). We investigated the differences in sequence composition of cPrDs from these two groups. The distribution of glutamine and asparagine content, a key driver of prion behavior (48), was similar in each (Fig. 7A, Supp. Fig. S10A). Yet in the sequences of the proteins that tested positive in both assays we observed an additional pattern: consecutive occurrence of proline and glutamine (P and Q) with nearby aromatic residues such as tyrosine and phenylalanine (Y and F; Supp. Fig. S10B,C). We observed no such signature in the proteins that tested negative in both assays (Supp. Fig. S10D). Tyrosine content was generally higher in the CR+/YPR+ group (Kruskal-Wallis H-test *P*=0.036; Wilcoxon rank-sum test *P*=0.042; Fig. 7A). The only CR+/YPR+ cPrD that contained no tyrosine, harbored the highest amount of phenylalanine (~5%) among all cPrDs. Tyrosine and phenylalanine content was significantly higher in the CR+/YPR+ group, and also in Q/N-rich PrDs of known prion proteins in yeast, than in the CR−/YPR− group (Kruskal-Wallis H-test *P*=0.01, 0.005; Wilcoxon rank-sum test *P*=0.01, 0.005, respectively; Fig. 7B). Moreover, the frequency of tyrosine and phenylalanine in PrDs of the only two currently known bacterial prion proteins – transcription termination factor Rho from *Clostridium botulinum* and single-stranded DNA-binding protein SSB from *Campylobacter hominis* – also revealed elevated levels as compared to our CR−/YPR− group (Fig. 7B). Thus, content of aromatic residues, especially of tyrosine and phenylalanine, in Q/N-rich PrDs may play a key role in determining the formation of prions across the evolutionary spectrum. Interestingly, a segment of the human prion protein (PrP) 169-YSNQNNF-175 containing glutamine, asparagine, tyrosine and phenylalanine, is required for its self-assembly (67). Moreover, the phenylalanine F175 is highly conserved, and the tyrosine Y169 is strictly conserved in mammalian PrPs and may functions as an aggregation gatekeeper (68) (Supp. Fig. S11). During self-assembly mediated by this segment, stacks of tyrosine and phenylalanine stabilize the amyloid core formed by the stacks of Q/Ns (69). This suggests role of tyrosine/phenylalanine in prion formation, not only limited to Q/N-rich PrDs.

**Fig. 7.**
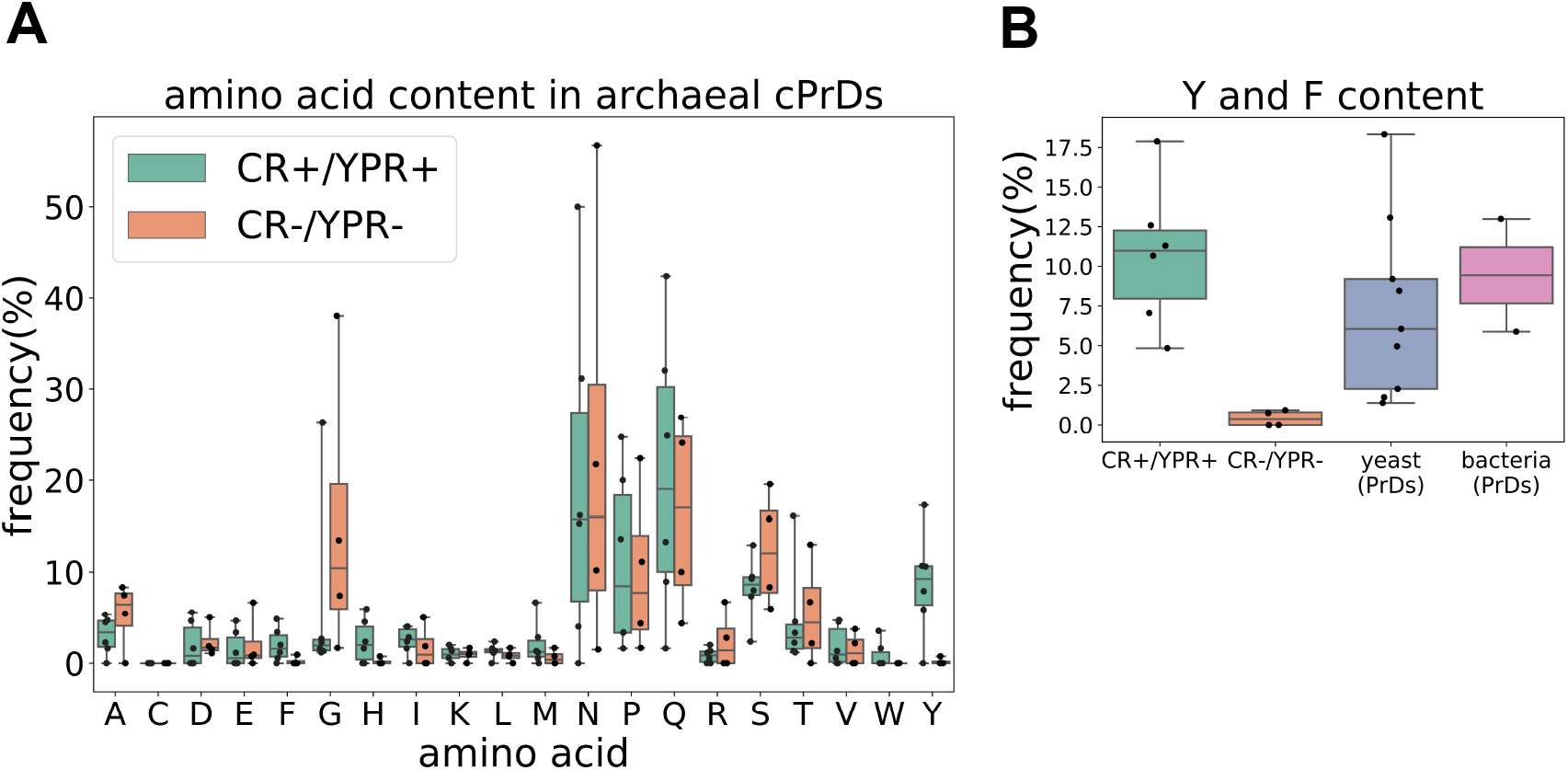
Tyrosine and/or phenylalanine distinguishes positive-testing cPrDs from negative-testing cPrDs. **A.** Comparison of amino acid frequency in cPrDs that tested positive in both Congo red and Sup35-based yeast-assay (CR+/YPR+), and ones that tested negative in both experimental assays (CR−/YPR−). Tyrosine (Y) content of the CR+/YPR+ cPrDs were higher than the CR−/YPR− cPrDs in general. The CR+/YPR+ cPrD with no tyrosine (lowest datapoint in distribution of Y) is the same with highest phenylalanine (F) frequency (highest data point in distribution of F). **B.** Frequencies of tyrosine and phenylalanine in archaeal cPrDs, yeast and bacterial PrDs. CR+/YPR+ cPrDs as well as yeast PrDs contain significantly higher Y/F than CR− /YPR− cPrDs. The Y/F frequency distribution in bacterial PrDs is also similar to that in CR+/YPR+ cPrDs and yeast PrDs. Since the bacterial group contained PrDs of only the two experimentally confirmed prions (Rho and SSB), no statistics were performed to compare the group with other ones.

The difference between CR−/YPR− and other groups was enhanced further when we compared the content of all amino acids with aromatic side chains (Supp. Fig. S10E). Thus, content of aromatic residues, especially of tyrosine and phenylalanine, in Q/N-rich PrDs may play a key role in determining the formation of prions across the evolutionary spectrum.

## Discussion

Long viewed as enigmatic drivers of disease, prions have emerged as a form of epigenetics beyond the chromosome that can be adaptive in eukaryotes ranging from fungi (24, 26, 27, 70, 71) to humans (72). Early analyses (48) suggested prions were absent in other domains of life, consistent with a prevailing view that they are evolutionarily young elements. The recent discovery of prions in bacteria (45) challenged this dogma, yet prions have not yet been identified in archaea. In this study we show that multiple archaeal cPrDs can form amyloids and act as *bona fide* elements of information transfer in living cells, fulfilling fundamental tenets of the prion characterization. To the best of our knowledge, neither prions nor intracellular amyloids have been reported in the domain Archaea to date. Confirming their presence would indicate their existence in all three domains of life, suggesting that prions were likely present during the time of LUCA.

Though many prions do not form amyloid (73–75), in the current study we focused on amyloid-forming cPrDs because recently identified bacterial prions have been shown to form amyloids (45), and amyloid fibril formation can also be detected with a relatively high-throughput experimental survey like the Congo red assay used here. We utilized the computational program PLAAC (49) to identify novel cPrDs. Although PLAAC’s underlying algorithm was trained on yeast PrDs, it has recently proven successful in facilitating the identification of prions in Bacteria (45, 76). Taken with our work here, in which PLAAC also enabled the identification of likely PrDs in Archaea, this speaks to the chemically conserved nature of at least a subset of (c)PrDs (amyloid-forming) across all three domains of life. We stress, however, that in this initial exploration of potential prions within the domain Archaea, we have performed a somewhat limited rather than exhaustive search (being guided by current information largely sourced from Q/N-rich, amyloid-forming, yeast-derived prions). Bacteria and archaeal proteomes appear to have fewer Q/N-rich regions in general as compared to eukaryotes (48), but it is as yet still unclear if this translates to them having fewer prions overall, or if it is the case that they are encoded with a different chemical signature to which we currently remain naive.

To investigate our computationally identified cPrDs experimentally, we utilized biochemical and genetic approaches (listed in regard to protein fragment, see Fig. 1). This included staining with amyloid specific dye (Fig. 4), infrared spectroscopy (Fig. 5A-C), transmission electron microscopy (Fig. 5D), and the yeast prion reporter assay (Fig. 6). We utilized the model organisms *E. coli* and *S. cerevisiae* in our experiments. Future work will require working with the source organism of the cPrD and testing the entire protein.

Looking across the archaeal domain, many species harbor at least 1 cPrD. However, Methanobacteria stand out as especially rich in cPrDs (Fig. 2, right side). Several members of the genera *Methanobrevibacter*, *Methanosphaera*, and *Methanobacterium* have five or more cPrDs per 1,000 proteins (Supp. Dataset 1). Out of 16 candidates tested, six were from the Class Methanobacteriaceae, two of which tested positive under both experimental assays (Table 1; Supp. Dataset 3). By far the dominant Gene Ontology (GO) term associated with the cPrDs identified in this group was integral component of membrane from the Cellular Component namespace (GO:0016021; ~70% of all assigned GO terms; Supp. Dataset 4), but this was also the case when looking exclusively outside of the Methanobacteria – with ~58% of all GO terms associated with cPrDs being GO:0016021, which was enriched overall (Fig. 3). The majority of the remaining cPrDs from the Methanobacteria were not annotated (Supp. Dataset 4). Possibly contributing to their relatively higher cPrD frequencies is that they had on average fewer proteins overall. The average number of proteins from the incorporated Methanobacteria was 1,915 ± 324 (mean ± 1SD), as compared to 2,238 ± 1,028 for all others. Likely further contributing is that the Q and N amino acids that comprise the Q/N-rich regions PLAAC is identifying are generally encoded by A/T-rich codons (Q = CAA or CAG; N = AAT or AAC; Supp. Dataset 5), and Methanobacteria have relatively higher AT-content compared to all others (65 ± 7.32% vs 51 ± 11.9%; Welch t.test p=7.64e-23). In general, those with higher AT content have higher normalized frequencies of cPrDs (Supp. Fig. S9).

Our GO enrichment analysis revealed that of the annotations that were significantly enriched or purified in proteins containing our cPrDs, the vast majority (~90%) were purified (e.g. depleted). As such, there is a high level of functional constraint with regard to which functional domains cPrDs tend to stably coexist with (meaning within the same protein). It is notable that overall metal ion binding (GO:0046872) is purified in the proteins harboring cPrDs relative to all proteins (Supp. Dataset 2), yet specific “child” GO terms nested underneath it such as calcium (GO:0005509) and copper ion binding (GO:0005507) are enriched (Fig. 3). This suggests there is something chemically consistent with Q/N-rich amyloids within these specific ion-binding proteins that is distinct from other metal ion-binding proteins. While we observed an enrichment in outer membrane cellular components and in several cell wall-related biological processes such as chitin binding, chitin metabolism, and cell wall adhesion (Fig. 3; Supp. Dataset 2), we note that it may simply be the case that cell-wall related proteins like these might tend to harbor Q/N-rich amyloids. This may place them disproportionately in the category of being detected by PLAAC, but not likely to present prion-like characteristics upon further experimental scrutiny. In this case, the only cell-wall related cPrD we tested was found to be negative in both the Congo red and yeast-prion reporter assays (cPrD2; adhesin-like protein; Table 1; Supp. Dataset 3).

The first prions discovered in the Domain Eukaryota (yeast translation release factor Sup35) (8) and the Domain Bacteria (transcription termination factor Rho) (45) were both related to cellular regulation. Further, amyloids in particular have been noted for their roles in signal transduction such as those involved with programmed cell death (77). In the current study, one of our positive-testing prion candidates was transcriptional repressor CopY (cPrD1; Table 1; Supp. Dataset 3), and one of our most highly enriched GO terms is involved with transcription regulation (GO:0003712; Fig. 3). As amyloid-based prions themselves can orchestrate a regulatory network (45, 72, 78, 79), this suggests that additional regulatory factors may be controlled by aggregation if they harbor a cPrD. This would provide a scenario in which the aggregation state of the protein (as dictated by intracellular conditions) might be a mechanism for controlling its activity. As in the case with the eukaryotic Sup35 (upon which aggregation results in less efficient translation termination) (18), and with the bacterial transcriptional terminator Rho (in which aggregation results in transcriptional read-through) (45), these types of “interferences” ultimately may provide a selective benefit by enabling a population under stress to rapidly explore a broader phenotypic landscape. Indeed, it has been demonstrated that prions in *Saccharomyces cerevisiae* can confer a fitness advantage (15, 17, 23). The observed association of cPrDs and confirmed PrDs being commonly found within genes responsible for cellular regulation across all three domains suggests PrDs may be an early-evolved manifestation of cellular organization that was likely present at the time of LUCA.

## Conclusion

With this work, we show that archaeal cPrDs can facilitate the acquisition of the prion phenotype, allowing them to function as protein-based elements of inheritance – thus expanding our knowledge of this epigenetic phenomenon to the third and final domain of life. This adds support to the hypothesis that amyloid-based prions were present at the earliest stages of life’s evolution. The current study represents an example of a top-down approach that pushes the story of amyloid-based functions further back in evolutionary time. This is complementary to work investigating the “amyloid world” theory of the origin of life (80, 81) that proposes that amyloids may have been forming from very short peptides before the emergence of what we might consider the first “living” system (82–86). These two approaches are heading toward each other, slowly closing the gap between them, with many interesting questions remaining to be answered and asked. Does a continuity exist between prebiotic amyloids and modern amyloid-based prions (87)? Are extant amyloid-based prions the biochemically evolved descendants of a prebiotic system capable of self-assembly and self-aggregation? Did the self-assembly and self-aggregation of these short peptides facilitate an expansion in range of potential interactions and functionalities? This work demonstrating protein-based, epigenetic inheritance via PrDs derived from the archaeal domain of life is a valuable step towards bridging this gap, but clearly there are many more to be taken.

## Materials and Methods

### Computational identification and classification of prion candidates

Archaeal reference proteomes and annotation information, including Gene Ontology (GO) annotations (51), were downloaded from the UniProt database (50). To systematically exclude low-quality proteomes, only those ranked as “standard” based on UniProt’s “Complete Proteome Detector” algorithm were incorporated, resulting in 1,262 proteomes holding 2,805,234 proteins as accessed on 27-Mar-2020.

The command-line version of PLAAC (49) (installed from https://github.com/whitehead/plaac in April of 2020) was used to identify proteins with putative prion domains in each proteome individually with default settings other than “-a 0” – which tells PLAAC to calculate and use the current proteome’s background amino acid frequencies entirely, rather than those of *Saccharomyces cerevisiae*. From those for which a core length of at least 60 was identified, several candidates that also possessed corresponding annotations were selected for experimental validation.

GO enrichment analyses were performed to identify enrichment/purification of frequencies of GO annotations in the proteins holding identified cPrDs (2,797) as compared to all scanned proteins (2,805,234) using the goatools v0.6.10 (52) “find_enrichment.py” script with default settings. Statistical significance was defined as those with Benjamini-Hochberg false-discovery rates of <=0.05. Many related GO terms were statistically significant at different depth levels and in those cases the term with the lowest depth was represented.

The phylogenomic tree was produced with GToTree v1.4.11 (88), within which 76 single-copy orthologs specific to the archaeal domain (using the “Archaea.hmm” included with GToTree) were identified with HMMER3 v3.2.1 (89), individually aligned with muscle v3.8.1551 (90), trimmed with trimal (91), and concatenated prior to phylogenetic estimation with FastTree2 v2.1.10 (92). Tree was initially visualized and edited with the Interactive Tree of Life interface (93).

Aggregation domains (20-amino-acid long) from prion candidates were selected for synthesis with pRANK (http://faculty.pieas.edu.pk/fayyaz/prank.html) (94) using default settings.

### Sequence analysis of experimentally tested archaeal cPrDs and known PrDs of yeast, and bacteria

We used glam2 from MEME suite version 5.1.1 (95, 96) for finding motifs via gapped alignment of CR+/YPR+ and CR−/YPR− cPrDs separately. For each group of cPrDs, keeping all other parameters set to default values, we programmatically performed runs of glam2 with varying values of the parameter “-n” which controls the number of iterations after which glam2 ends each run without improvement. For each glam2 run corresponding to a value of “n”, the number of alignment runs (“-r”, replicates) was set to 10 as a default. Initially the values of “n” were chosen from [4000, 6000, 8000, 10000, 20000, 40000, 60000, 80000, 100000]. At n=60000, we found the single optimal motif for the CR−/YPR− group to be reproduced by all 10 replicates. The score of this 30 amino acid long CR−/YPR− motif was 39.0692 (as shown in Supp. Figure S10D). On the other hand, although we found similar motifs across replicates for CR+/YPR+ cPrDs within the above-mentioned range of values for “n”, we strived to obtain identical motifs supported by multiple replicates. We then increased the number of iterations and chose the following values: [200000, 400000, 800000, 1000000, 2000000, 4000000]. At n=4million, we obtained the two optimal motifs with score 84.0115, and of length 42 and 40 amino acids, respectively (as shown in Supp. Figure S10B, S10C). Each of these motifs was supported by 5/10 replicates. They also showed similarity between themselves.

The coordinates of Q/N-rich prion domains (PrDs) of known yeast (*Saccharomyces cerevisiae*) prion proteins were obtained from Uniprot. The list of these proteins included: SWI1, SUP35, NEW1, RNQ1, MOT3, URE2, CYC8, SFP1, and NUP100. The PrD coordinates of NUP100 were obtained from *Halfmann et al. 2012* (97). The coordinates of bacterial PrD from transcription termination factor Rho (*Clostridium botulinum*) and single-stranded DNA-binding protein SSB (*Campylobacter hominis*) were obtained from Yuan and Hochschild 2017 and Fleming *et. al* 2019, respectively (45, 76). The PrD sequences were programmatically extracted from the full length fasta sequences of these proteins obtained from Uniprot.

The stats module from SciPy was used for the statistical tests.

### Colony color phenotype - Congo red assay

Curli-dependent amyloid generator (C-DAG) was used to direct the export of heterologous cPrDs to the cell surface of *E. coli* strain VS45 (53). Both bacterial strain and expression vector enabling protein production under the control of the arabinose-inducible PBAD promoter was part of comercially available C-DAG amylyloidogenicity kit https://www.kerafast.com/productgroup/540/c-dag-amyloidogenicity-kit. Export-directed fusion proteins contained the first 42 residues of CsgA signal sequence at the N terminus as described in the kit protocol. Sec translocon signal sequence is removed during translocation across the inner membrane, the 22-residue targeting sequence is retained at the N-terminus.

Lumio tag (CCPGCCGAGG) and His_6_ tag were added to the cPrD sequence at the C terminus during gene synthesis (www.idtdna.com)

For assessment of colony color phenotype, overnight cultures of VS45 were transformed with plasmids containing cPrDs. Bacteria were diluted to OD_600_ 0.01 in LB supplemented with the appropriate antibiotics (100 mg/mL carbenicillin, 25 mg/mL chloramphenicol). After 30 min of growth at 37 °C, 5 mL of the culture was spotted on LB agar plates supplemented with inducers (0.2% [w/v] L-arabinose, 1 mM IPTG), antibiotics (100 mg/mL carbenicillin; 25 mg/mL chloramphenicol), and, where indicated, on LB plates supplemented with Congo red (5 mg/ mL). Plates were then incubated for 4 days at room temperature. Photos of colonies were acquired using Canon EOS 6D SLR camera and the EOS Utility 3 software (https://www.usa.canon.com/internet/portal/us/home/support/self-help-center/eos-utility/).

### Light and fluorescent microscopy

Extracellular aggregates of heterologous candidate prion proteins stained with Congo Red were visualized using Zeiss Axioimager Z.1 with a Canon EOS SLR camera and 6D and EOS Utility 3 software.

### Transmission electron microscopy

Peptide solutions were adsorbed onto formvar/carbon-coated nickel grids (TED PELLA, CA, USA) in water, blotted dry, negatively stained with 1% uranyl acetate (TED PELLA, CA, USA), for 1 minute, blotted dry, and then viewed on transmission electron microscope Hitachi H-9500 TEM (Hitachi High Technologies America) at 300kV with an Advanced Microscopy Techniques 4-megapixel digital camera (AMT XR41B, Low-Dose CCD, USA).

### Peptide synthesis

Peptides corresponding to the 20 amino acid long fragments of putative PrDs were synthesized by ELIM Biopharm (Hayward, CA, USA). The peptides were of 95 % purity as determined by HPLC and mass spectrometry analyses. Peptides were dissolved in deionized water and stored at −80 °C. In our experiments, we used only the peptides that were initially soluble in water. pRANK-peptide sequences named P1, P2, P3, and P4 are presented in Fig. 5 and in the Supp. Dataset 3.

### Measurements of ThT fluorescence

For ThT fluorescence-based measurements, (λ ex. 440 nm/λ em. 485 nm) of fibrillization kinetics, a CLARIOstar plate reader from BMG LABTECH (Offenburg, Germany) and 96-well black microplates were used. All samples were slightly acidified to pH approximately 3. Typically, wells were filled with 150 μl volumes of diluted peptide samples containing ThT. Measurements were carried out at 37°C with agitation for 70 h; Each kinetic trace was calculated as an average from three independently collected trajectories (the error bars correspond to the standard deviations).

### ATR-FTIR

Measurements were carried out using single-reflection diamond ATR (attenuated total reflectance) accessory of the Nicolet iS50 FTIR spectrometer. The liquid suspensions of peptides were gently dried *in situ* until they formed films. Infrared spectra of the films were collected. Typically, for a single spectrum, 32 interferograms of 2 cm-1 resolution were co-added. Due to ambiguity in determining real values of refractive indices of amyloid aggregates, uncorrected ATR-FTIR data is shown. The spectra were corrected for water vapor, baseline and normalized. Data processing was performed using GRAMS software (Thermo Nicolet, USA).

### Yeast vector preparation

Fragments of genes corresponding to predicted prion domains were cloned into a yeast vector PDJ1776 (pAG415-ADH1-ccdB-SUP35C) using the Gateway Cloning enzymes (Invitrogen Clonase Gateway BP 11789020 and LR 11791020 Clonase II Enzyme Mix). Gateway Cloning procedures were based on methods described in Alberti et al. 2009 (46). Primers used to amplify cPrDs and to attach the recombinogenic attB1 and attB2 sites for Gateway cloning are listed in Supp. Dataset 6.

### Yeast strains

*Saccharomyces cerevisiae* strain YDJ1420 was transformed with plasmid pAG415-ADH1-cPrD#-SUP35C and subsequently cured of plasmid pAG426-GPD-SUP35C (plasmid shuffle). The resulting strains are labeled YDJ6455-6472. Example: strain YDJ6455-pAG415-ADH1-cPrD1-SUP35C (*Mata, leu2-3,112; his3-11,-15; trp1-1; ura3-1; ade1-14; can1-100; sup35::HygB; pAG415-ADH1-cPrD1-SUP35C [RNQ^+^], [psi^−^]*). *S. cerevisiae* strain YDJ1528 was used as positive control (*Mata, leu2-3,112; his3-11,-15; trp1-1; ura3-1; ade1-14; can1-100; sup35::HygB; AG415-ADH1-Sup35NM-Sup35C, [RNQ^+^], [psi^−^]*). *S. cerevisiae* strain YDJ1420 was used as negative control (*Mata, leu2-3,112; his3-11,-15; trp1-1; ura3-1; ade1-14; can1-100; sup35::HygB; pAG426-GPD-SUP35C, [RNQ^+^], [psi^+^]*). Standard rich (YPD) and Synthetic Dropout (SD) Media were used to culture yeast cells at 30 °C.

### Plasmid shuffling and phenotypic analysis

Plasmids containing archaeal prion domains fused with the carboxy-terminal domain of Sup35 (cPrD#-Sup35C) were, respectively, transformed into strain YDJ1420 by the standard transformation method (98). The resulting transformants were incubated on leucine-deficient synthetic medium (SD-Leu) for 3 days at 30 °C and then plated on SD-Leu medium supplemented with 1 mg/mL of 5-fluoroorotic acid (RPI Research Products F10501) to eliminate the Sup35 maintainer plasmid (pAG426-GPD-SUP35C) generating strains expressing an cPrD#-Sup35C fusion protein as the only source of functional Sup35. Phenotypes were assessed by the growth of colonies for 3 days at 30 °C in SD-Leu medium and plating of different dilutions of these cultures on YPD agar.

Post-shuffled yeast strains were plated on adenine-deficient synthetic medium (SD-Ade) medium to verify the readthrough status of the premature UGA stop codon in the *ade1-14* allele. In [*PSI*^+^] cells, the majority of Sup35 is aggregated unavailable for translation termination, which results in the read-through of the *ade1-14* premature stop codon. The synthesis of full-length Ade1 protein (N-succinyl-5-aminoimidazole-4-carboxamide ribotide synthetase) results in white colonies on rich medium and growth on SD-Ade medium. [*psi*^−^] cells showed no read-through of the *ade1-14* premature stop codon and did not grow on SD-Ade medium. Images of SD-Ade plates with colonies were analyzed with the ImageJ program (https://imagej.nih.gov/ij/) to quantitatively determine the outcome of the experiment. To be counted as a colony, the minimum area for a particle had to be 10% more than the largest particle area found in the negative control (area ≥ mm^2^) that has a circularity ≥ 0.75 (see Supp. Fig. 8.1, 8.2). The dish with at least one colony meeting these criteria is categorized as positive results of the yeast prion reporter assay (YPR+). If a dish has zero colonies, then it is categorized as a negative result (YPR−).

### Data and code availability

The code and data for reproducing the computational analyses performed herein are available at https://figshare.com/projects/Zajkowski_et_al_2020_Archaeal_prion_data_and_code_repository/78720.

## Supporting information

PLAAC results per genome

GO enrichment analysis results

Experimentally tested cPrD list and information

All PLAAC positive proteins and annotations

Amino-acid codon GC and AT content

Primers used in preparation of cPrDs for Gateway cloning

## Acknowledgments

- Tomasz Zajkowski was funded by Ministerstwo Nauki i Szkolnictwa Wyższego under the Mobilność Plus program (1656/MOB/V/2017/0).
- Amanda Carbajal was funded by M058903-20 NIH/NIGMS grant under the IMSD program.
- TEM image acquisition made possible by the support of the Universities Space Research Association (USRA) Science & Technology Innovation Labs Program managed under NASA Contract NNA16BD14C
- We thank Thomas Lozanoski for his help in the yeast prion reporter assays.
- We thank Anupam Chakravarty for his advice on the experimental approach.
- We thank Stanisław Dunin-Horkawicz for his intellectual support at the early stages of the project.
- We acknowledge the support of the Origins Research Group of Exobiology Laboratory (ORGEL) at NASA Ames Research Center

