## Supplementary figures for "The hunt for ancient prions: Archaeal prion-like domains form amyloids and substitute for yeast prion domains"

**This PDF file includes:**

Figures S1 to S11
Legends for Datasets S1 to Dataset S6

**Other supplementary materials for this manuscript include the following:**

Datasets S1 to Dataset S6
Data and code availability

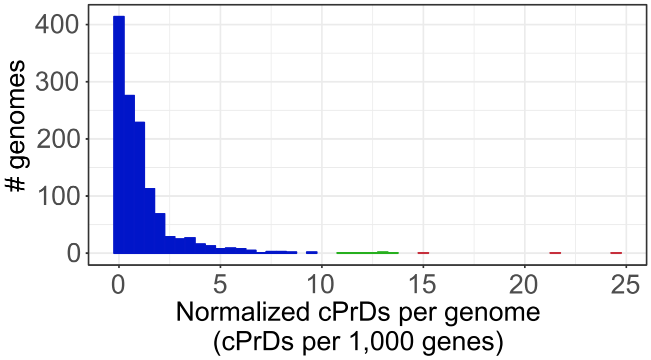


**Fig. S1.** Histogram depicting the distribution of normalized PLAAC-identified cPrD counts per genome (N = 1,262). Green are those >= 10, red are >=15.

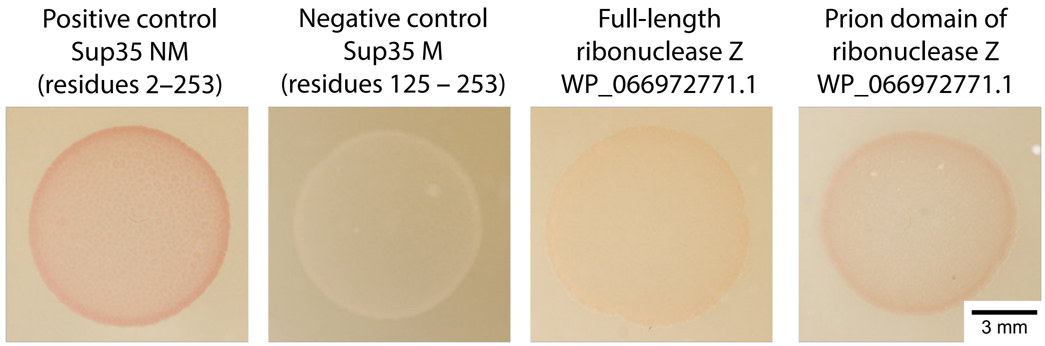


**Fig. S2.** Bacterial colonies expressing cPrD have stronger color than bacterial colonies expressing the full-length protein. Comparison of color intensity of bacterial colonies exporting full-length ribonuclease Z (WP_066972771.1) from *Methanobrevibacter filiformis* and it’s cPrD growing on media supplemented with 0.1% Congo red.

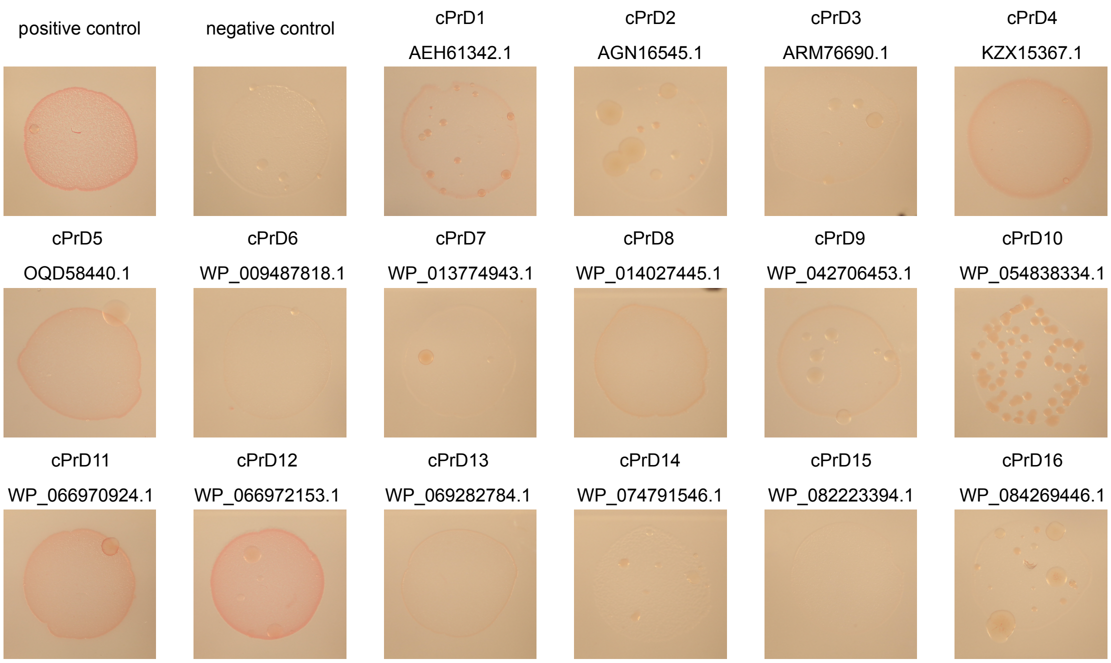


**Fig. S3.** Colonies of *E. coli* grew on the induction medium containing 0.1% Congo red. The red color of the colony indicates that the expressed protein binds to Congo red. Binding of Congo red is typical for amyloid fibrils. Sup35 NM (residues 2-253) used as a positive control. Sup35 M (residues 125-253) used as a negative control.

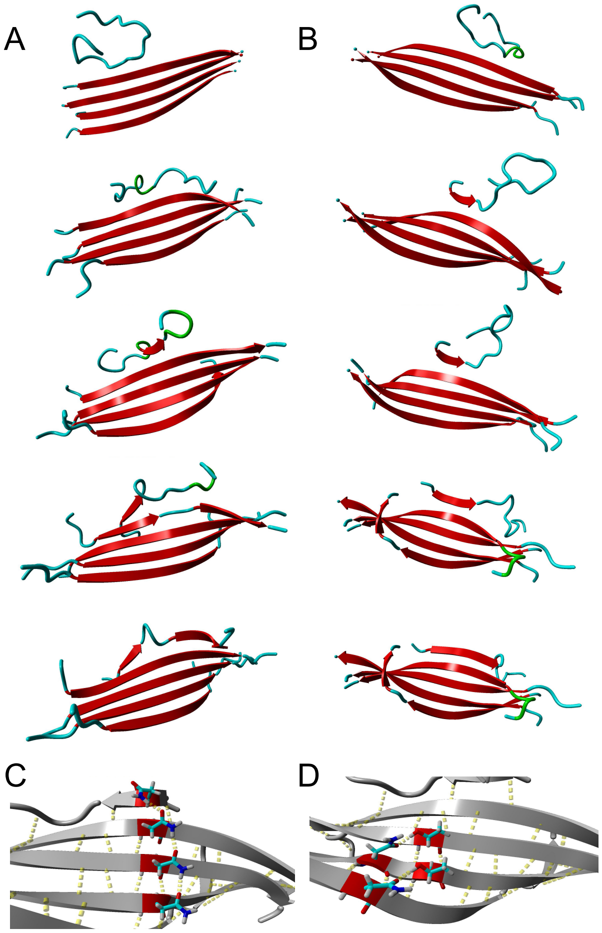


**Fig. S4.** Molecular dynamics of peptide P2 (KPQNNTNNTTNNNTTNNNTN) showing that presence of β-sheet P2 (red-parallel) promotes the formation of a new filament of β-sheet out of unfolded P2 peptide (consecutive timepoints from top to bottom). A. formation of parallel β-sheet filament over 300 ns of simulation and B. formation of nonparallel sheet over 200 ns of simulation. Molecular dynamics was performed on Yasara 19.9.17 under Amber 03 force-field with water as solvent and periodic boundary conditions. The ease of promotion of β-sheet might be a reason of additional structure stabilization by Asn and Thr residues interactions which may form C. a perpendicular (in comparison to the β-sheet filaments) chain of hydrogen bonds formed between Asn residues or/and D. pairs of hydrogen bonds between Asn and Thr across the neighboring filaments. Such interactions of side chains of amino acids form additional two plains of hydrogen bonds (before and behind the β-sheet plane) which may significantly increase the stability of such a secondary structure of a peptide/protein. Although the promotion of β-sheet formation most probably occurs in similar sequences when exposed on formed β-sheet the position of the newly formed filament seems to be random, therefore it most probably might need other structures to position it within the amyloid.

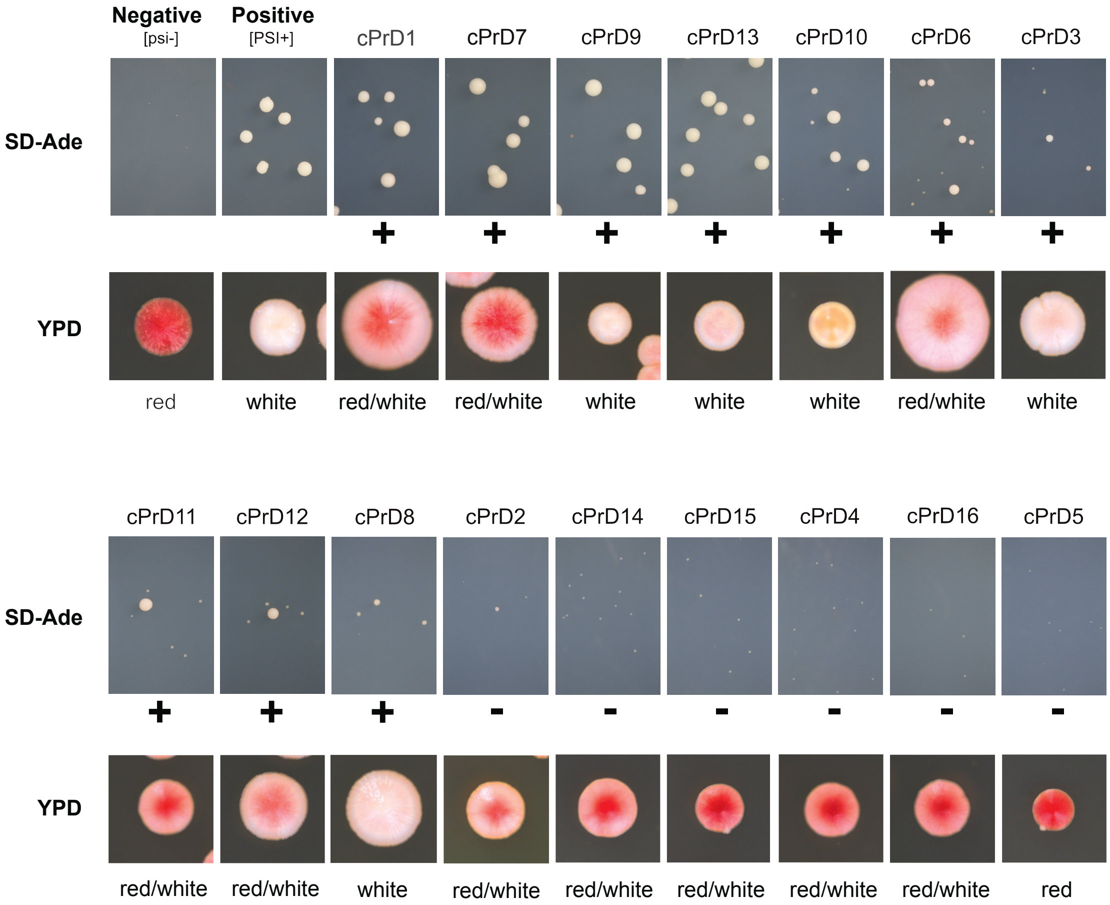


**Fig. S5.** cPrD-SUP35C strains display different ability to grow on media lacking adenine and a variety of colony colors. Images of representative colonies of cPrD-Sup35C-expressing yeast strains growing on SD-Ade and YPD plates. Colonies phenotypes [*psi*-] and [*PSI*+] are shown for comparison. Positive results are marked with “+” and are considered positive results of the yeast prion reporter assay (YPR+). Negative results are marked with “-” and are considered negative results of the yeast prion.

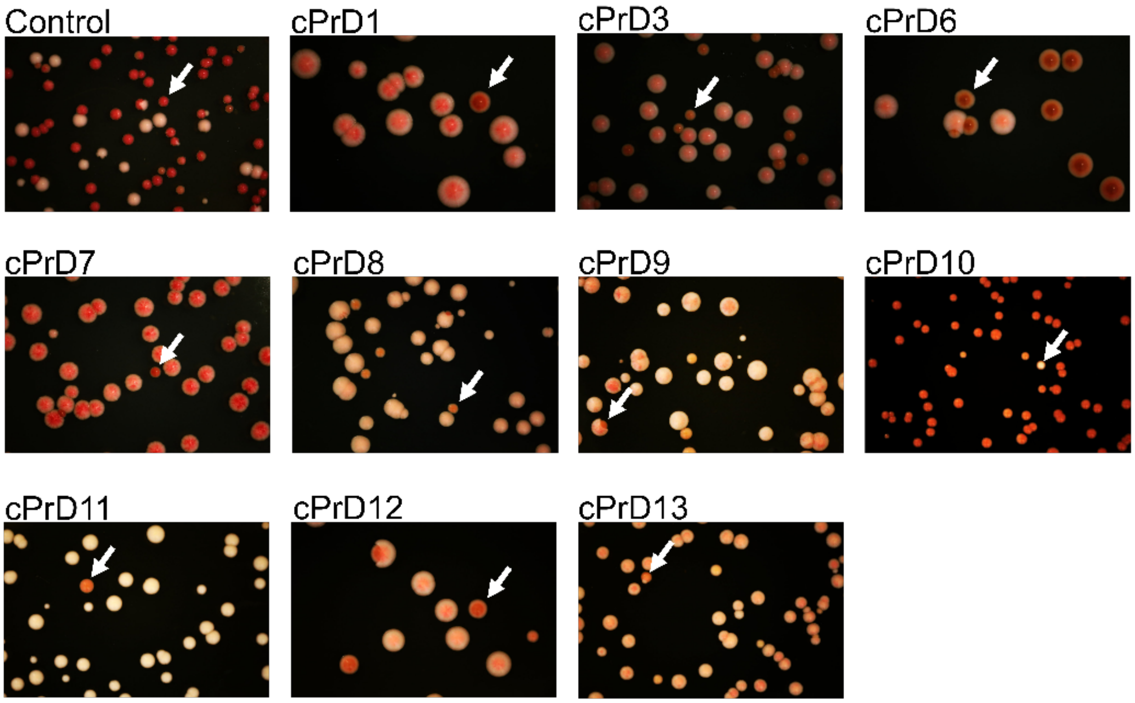


**Fig. S6.** Reversion of prion phenotype. Colonies harboring cPrD-Sup35MC that generated a phenotype resembling *[PSI+]* (YPR+ from Supp. Fig. S5) were restreaked on YPD media. Some colonies show reversion to *[psi-]* (revealed by red coloration) phenotype suggesting that cPrD-Sup35MC is in a metastable state allowing reversion of prion phenotype. Arrows mark colonies of reversed phenotype. In the case of cPrD10 the only colony that did not reverse is marked. Reversion of *[psi-]* phenotype of colonies harboring Sup35 full-length protein shown as a control.

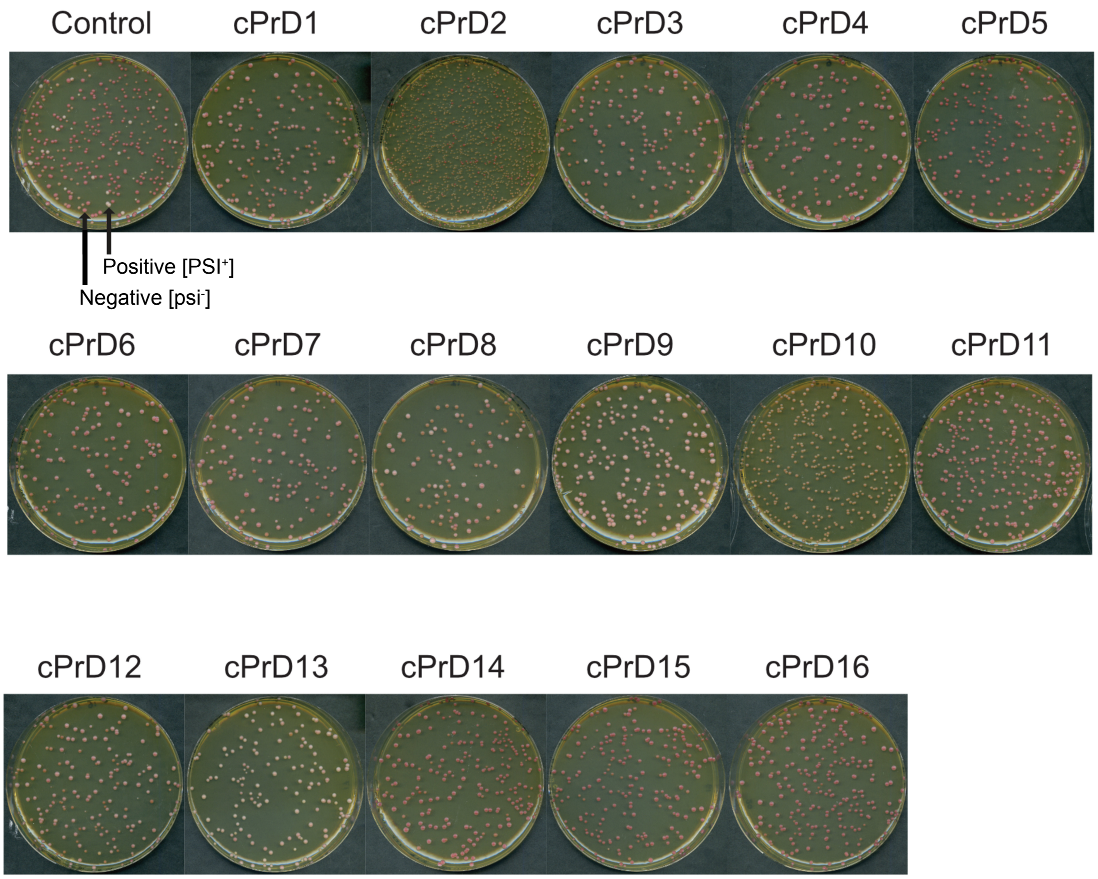


**Fig. S7.** Images of the colonies of 16 cPrD-Sup35C-expressing strains growing on YPD media. A strain with full-length Sup35 was used as control.

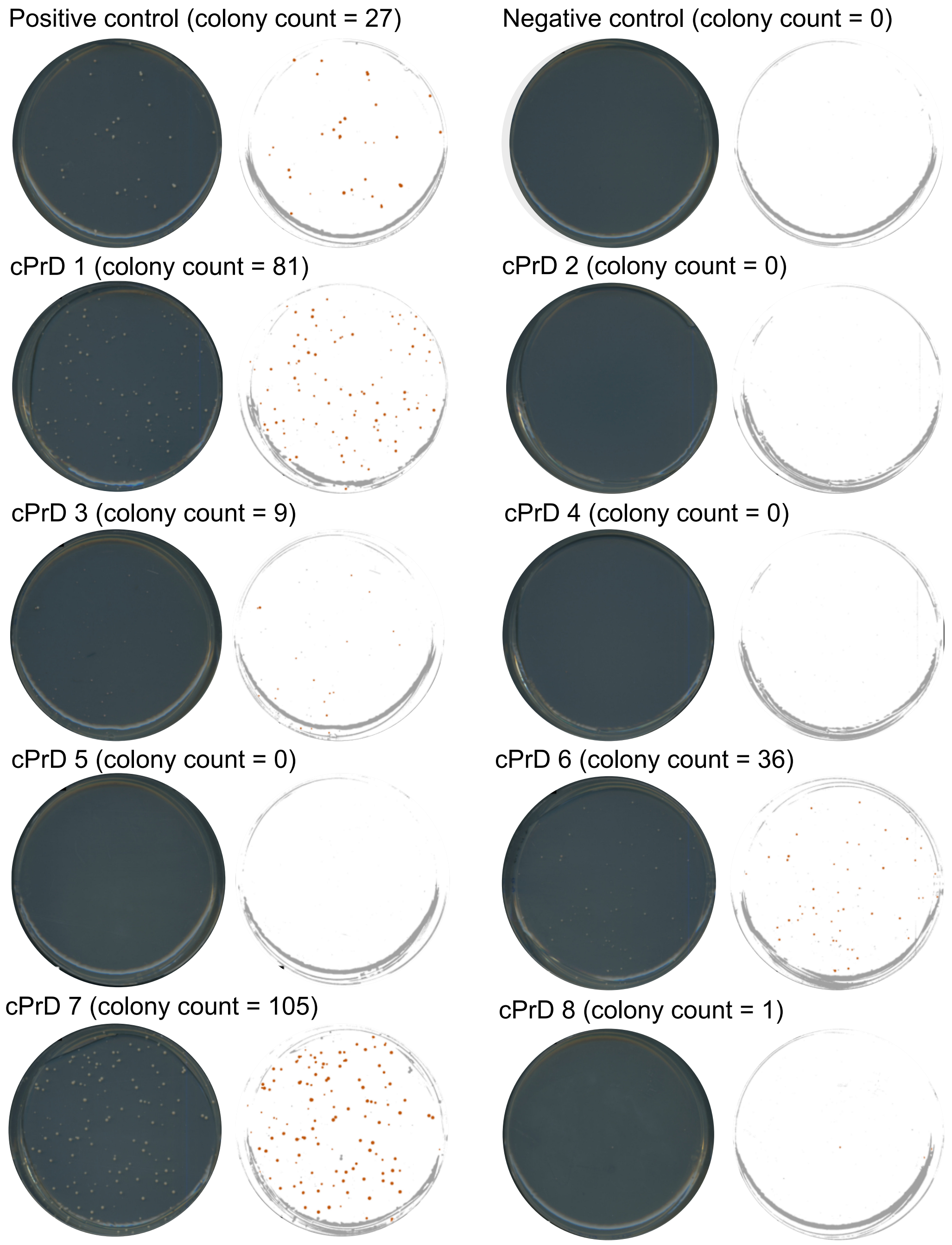


**Fig. S8.1** Petri dishes (9 cm diameter) with cPrD-Sup35C-expressing strains on SD-Ade. Section A: Photo of Petri dish. Section B: Image analyzed with ImageJ (https://imagej.nih.gov/ij/). Particles with an area > 0.35 $mm^2$ and a roundness > 0.75 are marked in red and counted as colonies. Particles not fulfilling these criteria are marked in gray. Positive control strain with full-length Sup35. Negative control strain with the C-terminal part of Sup35.

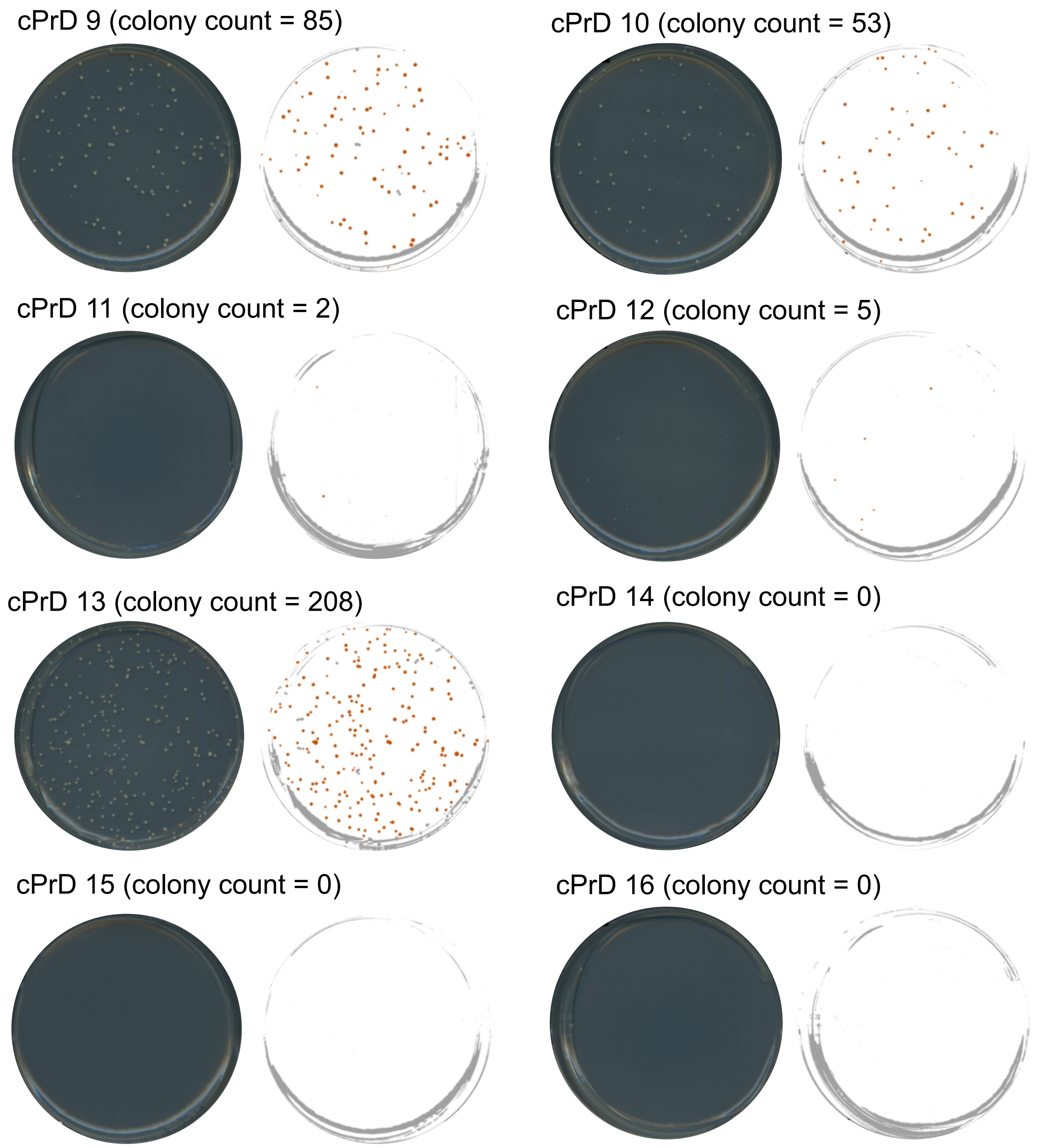


**Fig. S8.2** Petri dishes (9 cm diameter) with cPrD-Sup35C-expressing strains on SD-Ade. Section A: Photo of Petri dish. Section B: Image analyzed with ImageJ (https://imagej.nih.gov/ij/). Particles with an area > 0.35 $mm^2$ and a roundness > 0.75 are marked in red and counted as colonies. Particles not fulfilling these criteria are marked in gray. Positive control strain with full-length Sup35. Negative control strain with the C-terminal part of Sup35.

**Normalized cPrDs as a function of AT content**

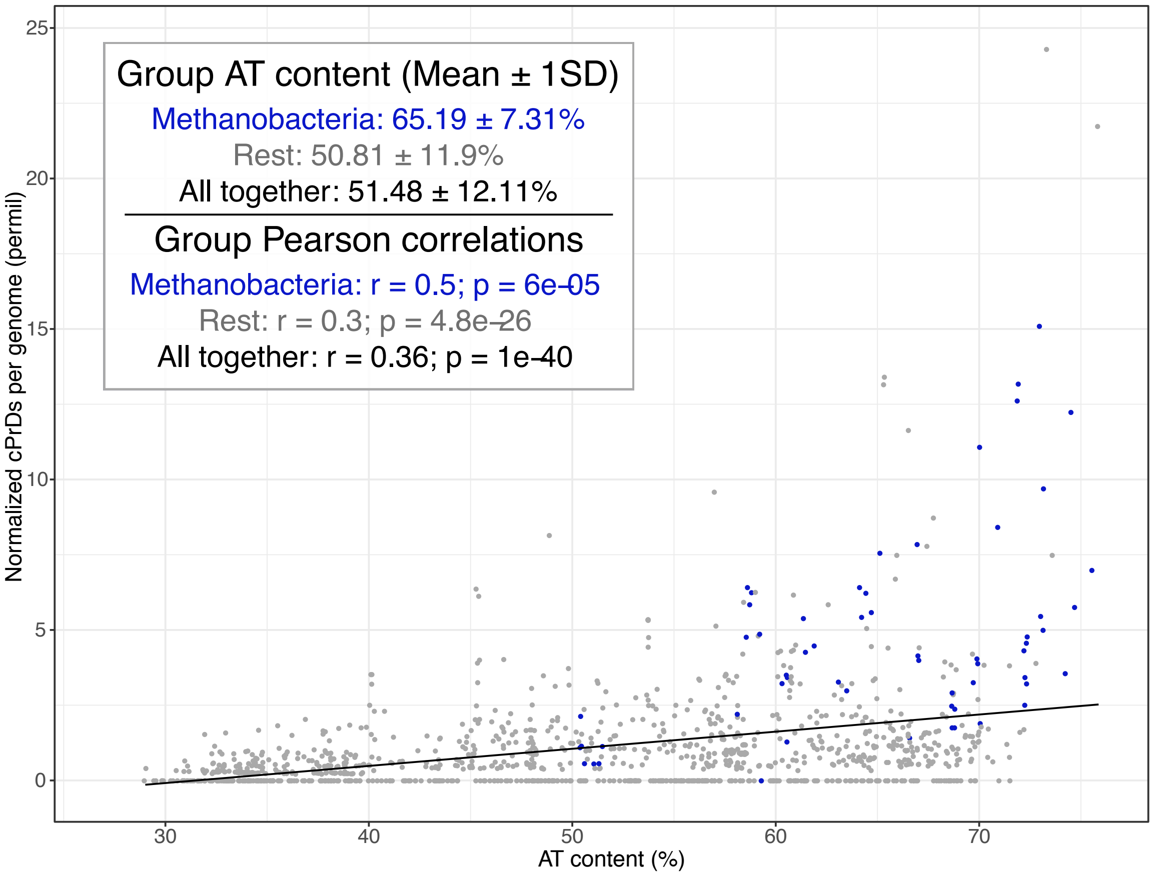


**Fig. S9**. Normalized cPrDs plotted as a function of AT content for each of the incorporated genomes (N = 1,262). Representatives of the Methanobacteria class are colored blue.

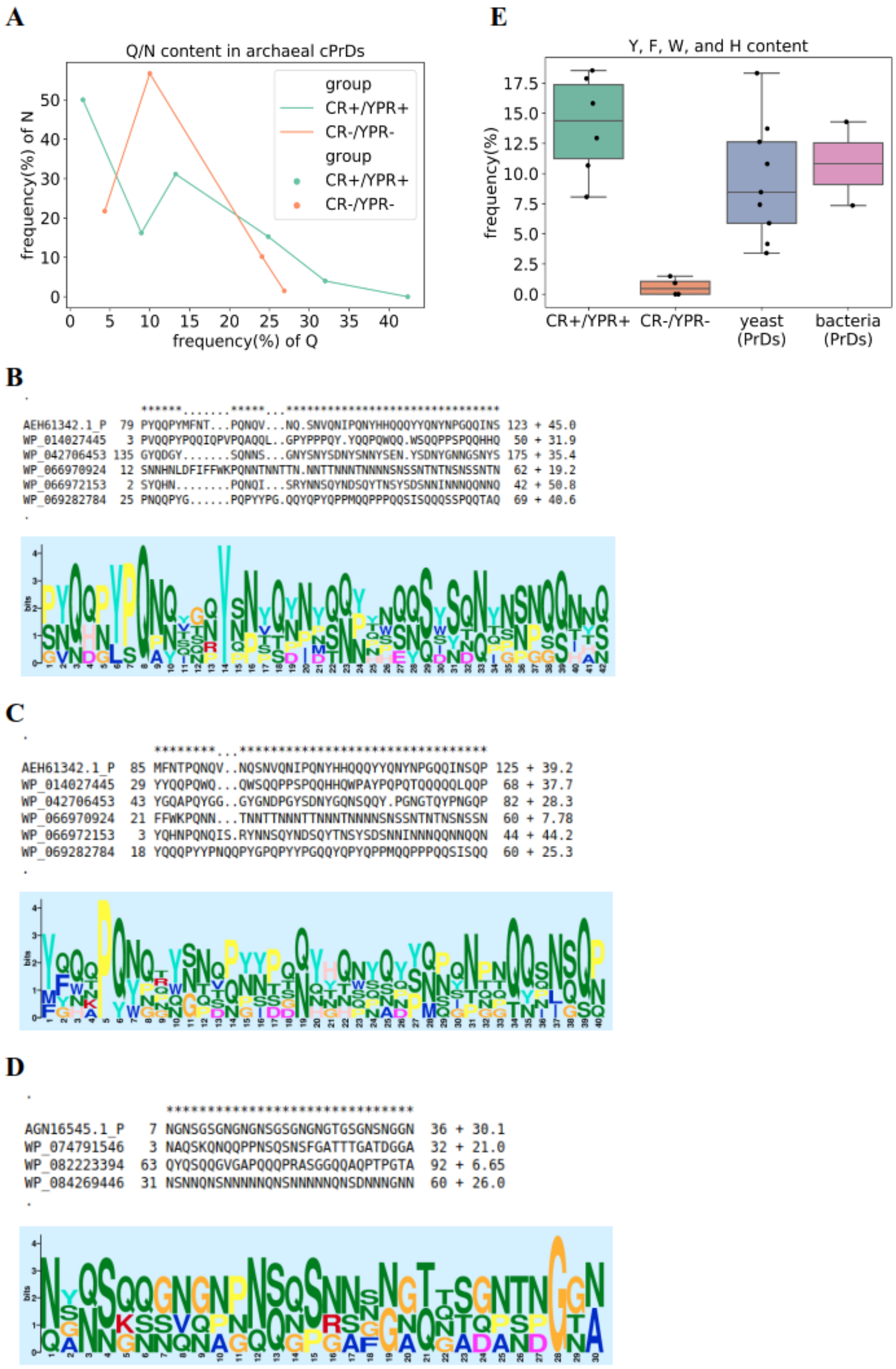
A
Q/N content in archaeal cPrDs
frequency(%) of N
frequency(%) of Q
group
CR+/YPR+
CR-/YPR-
E
Y, F, W, and H content
frequency(%)
CR+/YPR+
CR-/YPR-
yeast (PrDs)
bacteria (PrDs)
B
******.......*****...******************************
AEH61342.1_P 79 PYQQPYMFNT...PQNQV...NQ.SNVQNIPQNYHHQQQYYQNYNPGQQINS 123 + 45.0
WP_014027445 3 PVQQPYPQQIQPVPQAQQL..GPYPPPQY.YQQPQWQQ.WSQQPPSPQQHHQ 50 + 31.9
WP_042706453 135 GYQDGY.......SQNNS...GNYSNYSDNYSNNYSEN.YSDNYGNNGSNYS 175 + 35.4
WP_066970924 12 SNNHNLDFIFFWKPQNNTNNTTN.NNTTNNNTNNNNSNSSNTNTNSNSSNTN 62 + 19.2
WP_066972153 2 SYQHN........PQNQI...SRYNNSQYNDSQYTNSYSDSNNINNNQQNNQ 42 + 50.8
WP_069282784 25 PNQQPYG......PQPYYPG.QQYQPYQPPMQQPPPQQSISQQQSSPQQTAQ 69 + 40.6
C
********...*********************************
AEH61342.1_P 85 MFNTPQNQV..NQSNVQNIPQNYHHQQQYYQNYNPGQQINSQP 125 + 39.2
WP_014027445 29 YYQQPQWQ...QWSQQPPSPQQHHQWPAYPQPQTQQQQQLQQP 68 + 37.7
WP_042706453 43 YGQAPQYGG..GYGNDPGYSDNYGQNSQQY.PGNGTQYPNGQP 82 + 28.3
WP_066970924 21 FFWKPQNN...TNNTTNNNTTNNNTNNNNSNSSNTNTNSNSSN 60 + 7.78
WP_066972153 3 YQHNPQNQIS.RYNNSQYNDSQYTNSYSDSNNINNNQQNNQQN 44 + 44.2
WP_069282784 18 YQQQPYYYPNQQPYGPQPYYPGQQYQPYQPPMQQPPPQQSISQQ 60 + 25.3
D
******************************
AGN16545.1_P 7 NGNSGSGNGNGNSGSGNGNGTGSGNSNGGN 36 + 30.1
WP_074791546 3 NAQSKQNQQPPNSQSNSFGATTTGATDGGA 32 + 21.0
WP_082223394 63 QYQSQQGVGAPQQQPRASGGQQAQPTPGTA 92 + 6.65
WP_084269446 31 NSNNQNSNNNNNQNSNNNNNQNSDNNNGNN 60 + 26.0

**Fig. S10.** Sequence features of positive-testing and negative-testing cPrDs. (A) Q/N content in archaeal cPrDs. The frequency of Q and N in each cPrD were plotted for both CR+/YPR+ and CR-/YPR- groups. Sequence alignment and logo of sequence motifs found with glam2 for CR+/YPR+ group (B, C for motif 1 and motif 2, respectively) and CR-/YPR- group (D). The pattern of consecutive proline and glutamine flanked by tyrosine/phenylalanine or other aromatic amino acids was observed in the motifs from CR+/YPR+ group, but not in the motif from CR-/YPR- group. (E) Frequencies of amino acid residues with aromatic side-chains (tyrosine, phenylalanine, tryptophan and histidine) in archaeal cPrDs, yeast and bacterial PrDs. CR+/YPR+ cPrDs as well as yeast PrDs contain significantly higher aromatic amino acid residues than CR-/YPR- cPrDs (t-test *P*= 0.0002, 0.0049; Kruskal-Wallis H-test *P*= 0.01, 0.005; Wilcoxon rank-sum test *P*= 0.01, 0.005, respectively for CR+/YPR+ vs CR-/YPR-, and yeast PrDs vs CR-/YPR-). The frequency distribution in bacterial PrDs also revealed elevated levels compared to that in CR-/YPR- group. Since the bacterial group contained PrDs of only the two experimentally confirmed prions (Rho and SSB), no statistics were performed to compare the group with the other groups.

```
P10279 PRIO_BOVIN   173   YYRPVDQYSNQNNFVHDCVNITVKEHTVTTTTKGENFTETDIKMMERVVEQMCITQYQRE   232
P04156 PRIO_HUMAN   162   YYRPMDEYSNQNNFVHDCVNITIKQHTVTTTTKGENFTETDVKMMERVVEQMCITQYERE   221
P04925 PRIO_MOUSE   161   YYRPVDQYSNQNNFVHDCVNITIKQHTVTTTTKGENFTETDVKMMERVVEQMCVTQYQKE   220
B5SY89 PRIO_BOBOX   173   YYRPVDQYSNQNNFVHDCVNITVKEHTVTTTTKGENFTETDIKMMERVVEQMCITQYQRE   232
P04273 PRIO_MESAU   162   YYRPVDQYNNQNNFVHDCVNITIKQHTVTTTTKGENFTETDIKIMERVVEQMCTTQYQKE   221
P23907 PRIO_SHEEP   165   YYRPVDRYSNQNNFVHDCVNITVKQHTVTTTTKGENFTETDIKIMERVVEQMCITQYQRE   224
P13852 PRIO_RAT     162   YYRPVDQYSNQNNFVHDCVNITIKQHTVTTTTKGENFTETDVKMMERVVEQMCVTQYQKE   221
Q95211 PRIO_RABIT   161   YYRPVDQYSNQNSFVHDCVNITVKQHTVTTTTKGENFTETDIKIMERVVEQMCITQYQQE   220
P49927 PRIO_PIG     166   YYRPVDQYSNQNSFVHDCVNITVKQHTVTTTTKGENFTETDVKMIERVVEQMCITQYQKE   225
P61768 PRIO_PANTR   162   YYRPMDQYSSQNNFVHDCVNITIKQHTVTTTTKGENFTETDVKMMERVVEQMCITQYERE   221
P40244 PRIO_NEOVI   166   YYKPVDQYSNQNNFVHDCVNITVKQHTVTTTTKGENFTETDMKIMERVVEQMCVTQYQRE   225
P67992 PRIO_MACFA   162   YYRPVDQYSNQNNFVHDCVNITIKQHTVTTTTKGENFTETDVKMMERVVEQMCITQYEKE   221
P40252 PRIO_GORGO   162   YYRPMDQYSNQNNFVHDCVNITIKQHTVTTTTKGENFTETDVKMMERVVEQMCITQYERE   221
P79141 PRIO_CAMDR   165   YYKPVDQYSNQNSFVHDCVNITVKQHTVTTTTKGENFTETDVKMMERVVEQMCITQYQRE   224
Q60506 PRIO_CRIGR   162   YYRPVDQYNNQNNFVHDCVNITIKQHTVTTTTKGENFTETDVKMMERVVEQMCVTQYQKE   221
P67997 PRIO_MACMU   162   YYRPVDQYSNQNNFVHDCVNITIKQHTVTTTTKGENFTETDVKMMERVVEQMCITQYEKE   221
O18754 PRIO_FELCA   165   YYRPVDQYSNQNNFVHDCVNITVKQHTVTTTTKGENFTETDMKIMERVVEQMCVTQYQKE   224
P52113 PRIO_CAPHI   165   YYRPVDQYSNQNNFVHDCVNITVKQHTVTTTTKGENFTETDIKIMERVVEQMCITQYQRE   224
P67986 PRIO_CEREN   165   YYRPVDQYNNQNTFVHDCVNITVKQHTVTTTTKGENFTETDIKMMERVVEQMCITQYQRE   224
P40247 PRIO_CALJA   161   YYRPVDQYNNQNNFVHDCVNITIKQHTVTTTTKGENFTETDVKMMERVVEQMCITQYEKE   220
P52114 PRIO_MUSPF   166   YYKPVDQYSNQNNLVHDCVNITVKQHTVTTTTKGENFTETDMKIMERVVEQMCVTQYQQE   225
P40251 PRIO_COLGU   162   YYRPVDQYSNQNNFVHDCVNITIKQHTVTTTTKGENFTETDVKMMERVVEQMCITQYEKE   221
Q5UJI7 PRIO_BOSIN   173   YYRPVDQYSNQNNFVHDCVNITVKEHTVTTTTKGENFTETDIKMMERVVEQMCITQYQRE   232
P67993 PRIO_MACAR   162   YYRPVDQYSNQNNFVHDCVNITIKQHTVTTTTKGENFTETDVKMMERVVEQMCITQYEKE   221
P40255 PRIO_MANSP   155   YYRPVDQYSNQNNFVHDCVNITIKQHTVTTTTKGENFTETDVKMMERVVEQMCITQYEKE   214
P40257 PRIO_TRAFR   162   YYRPVDQYSNQNNFVHDCVNITIKQHTVTTTTKGENFTETDVKMMERVVEQMCITQYEKE   221
P67989 PRIO_CERDI   154   YYRPVDQYSNQNNFVHDCVNITIKQHTVTTTTKGENFTETDVKMMERVVEQMCITQYEKE   213
P67995 PRIO_MACNE   162   YYRPVDQYSNQNNFVHDCVNITIKQHTVTTTTKGENFTETDVKMMERVVEQMCITQYEKE   221
P40258 PRIO_SAISC   169   YYRPVDQYSNQNNFVHDCVNVTIKQHTVTTTTKGENFTETDVKMMERVVEQMCITQYEKE   228
Q95270 PRIO_THEGE   147   YYRPVDQYSNQNNFVHDCVNITIKQHTVTTTTKGENFTETDVKMMERVVEQMCITQYQKE   206
P67990 PRIO_LOPAT   147   YYRPVDQYSNQNNFVHDCVNITIKQHTVTTTTKGENFTETDVKMMERVVEQMCITQYEKE   206
P40245 PRIO_AOTTR   154   YYRPVDQYSNQNNFVHDCVNITIKQHTVTTTTKGENFTETDVKIMERVVEQMCITQYEKE   213
P67988 PRIO_CHLAE   154   YYRPVDQYSNQNNFVHDCVNITIKQHTVTTTTKGENFTETDVKMMERVVEQMCITQYEKE   213
P40256 PRIO_PONPY   162   YYRPVDQYSNQNNFVHDCVNITIKQHTVTTTTKGENFTETDVKMMERVVEQMCITQYERE   221
Q95176 PRIO_CERAT   155   YYRPVDQYSNQNNFVHDCVNITIKQHTVTTTTKGENFTETDVKMMERVVEQMCITQYEKE   214
P67996 PRIO_PAPHA   162   YYRPVDQYSNQNNFVHDCVNITIKQHTVTTTTKGENFTETDVKMMERVVEQMCITQYEKE   221
P67994 PRIO_MACFU   162   YYRPVDQYSNQNNFVHDCVNITIKQHTVTTTTKGENFTETDVKMMERVVEQMCITQYEKE   221
P40249 PRIO_SAPAP   161   YYRPVDQYSNQNNFVHDCVNITIKQHTVTTTTKGENFTETDVKMMERVVEQMCITQYERE   220
P61761 PRIO_CERMO   155   YYRPVDQYSNQNNFVHDCVNITIKQHTVTTTTKGENFTETDVKMMERVVEQMCITQYEKE   214
P61766 PRIO_HYLLA   162   YYRPMDQYSSQNNFVHDCVNITIKQHTVTTTTKGENFTETDVKMMERVVEQMCITQYERE   221
P40246 PRIO_ATEGE   146   YYRPVDQYNNQNNFVHDCVNITIKQHTVTTTTKGENFTETDVKMMERVVEQMCITQYERE   205
P61767 PRIO_SYMSY   162   YYRPMDQYSSQNNFVHDCVNITIKQHTVTTTTKGENFTETDVKMMERVVEQMCITQYERE   221
Q95174 PRIO_ERYPA   155   YYRPVDQYSNQNNFVHDCVNITIKQHTVTTTTKGENFTETDVKMMERVVEQMCITQYEKE   214
P40248 PRIO_PLEMO   155   YYRPVDQYSNQNNFVHDCVNITIKQHTVTTTTKGENFTETDVKMMERVVEQMCITQYEKE   214
P61762 PRIO_CERNE   155   YYRPVDQYSNQNNFVHDCVNITIKQHTVTTTTKGENFTETDVKMMERVVEQMCITQYEKE   214
P67991 PRIO_MACSY   147   YYRPVDQYSNQNNFVHDCVNITIKQHTVTTTTKGENFTETDVKMMERVVEQMCITQYEKE   206
P51446 PRIO_ATEPA   161   YYRPVDQYNNQNNFVHDCVNITIKQHTVTTTTKGENLTETDVKMMERVVEQMCITQYERE   220
Q60468 PRIO_CRIMI   162   YYRPVDQYNNQNNFVHDCVNITIKQHTVTTTTKGENFTETDVKMMERVVEQMCVTQYQKE   221
O46501 PRIO_CANLF   165   YYRSVDQYNNQSTFVHDCVNITVKQHTVT-TTKGENFTETDIKMMERVVEQMCITQYQRE   223
P47852 PRIO_ODOHE   165   YYRPVDQYNNQNTFVHDCVNITVKQHTVTTTTKGENFTETDIKMMERVVEQMCITQYQRE   224
Q6EH52 PRIO_AILME   173   YYRPVDQYSNQNNFVHDCVNITVKEHTVTTTTKGENFTETDIKMMERVVEQMCITQYQRE   232
Q5UJG7 PRIO_BOSTR   173   YYRPVDQYSNQNNFVRDCVNITVKEHTVTTTTKGENFTETDIKMMERVVEQMCITQYQTE   232
Q5UAF1 PRIO_BISBI   173   YYRPVDQYSNQNNFVHDCVNITVKEHTVTTTTKGENFTETDIKMMERVVEQMCITQYQRE   232
Q5UJH0 PRIO_BOSGA   173   YYRPVDQYSNQNNFVHDCVNITVKEHTVTTTTKGENFTETDIKMMERVVEQMCITQYQRE   232
Q5UJG3 PRIO_TRAIM   173   YYRPVDQYSNQNNFVHDCVNITVKQHTVTTTTKGENFTETDIKMMERVVEQMCITQYQRE   232
Q5UJH8 PRIO_BUBBU   173   YYRPVDQYSNQNNFVHDCVNITVKEHTVTTTTKGENFTETDIKMMERVVEQMCITQYQRE   232
Q7JK02 PRIO_OVIMU   165   YYRPVDQYSNQNNFVHDCVNITVKQHTVTTTTKGENFTETDIKIMERVVEQMCITQYQRE   224
Q5UJG1 PRIO_ANTCE   173   YYRPVDQYSNQNNFVHDCVNITVKQHTVTTTTKGENFTETDIKMMERVVEQMCITQYQRE   232
Q95M08 PRIO_BUDTA   165   YYRPVDQYSNQNNFVHDCVNITVKQHTVTTTTKGENFTETDIKIMERVVEQMCITQYQRE   224
Q7JIY2 PRIO_OVIMO   165   YYRPVDQYSNQNNFVHDCVNITVKQHTVTTTTKGENFTETDIKIMERVVEQMCITQYQRE   224
Q9Z0T3 PRIO_SIGHI   162   YYRPVDQYNNQNNFVHDCVNITIKQHTVTTTTKGENFTETDVKMMERVVEQMCVTQYQKE   221
Q68G95 PRIO_MOSCH   165   YYRPVDQYSNQNNFVHDCVNITVKQHTVTTTTKGENFTETDIKIMERVVEQMCITQYQRE   224
P67987 PRIO_CEREL   165   YYRPVDQYNNQNTFVHDCVNITVKQHTVTTTTKGENFTETDIKMMERVVEQMCITQYQRE   224
Q7JIH3 PRIO_OVICA   165   YYRPVDQYSNQNNFVHDCVNITVKQHTVTTTTKGENFTETDIKIMERVVEQMCITQYQRE   224
Q5XVM4 PRIO_RUPRU   165   YYRPVDQYSNQNNFVHDCVNITVKQHTVTTTTKGENFTETDIKIMERVVEQMCITQYQRE   224
B0FYL5 PRIO_VULLA   166   YYRPVDQYSNQNNFVRDCVNITVKQHTVTTTTKGENFTETDMKIMERVVEQMCVTQYQKE   225
P51780 PRIO_TRIVU   167   MYRPIDQYSSQNNFVHDCVNITVKQHTTTTTTKGENFTETDIKIMERVVEQMCITQYQAE   226
                           *: :*.*..*..:*:****:*:*:**.* ******:****:*::******** ***: *
```

**Fig. S11.** Sequence alignment of mammalian prion proteins (PrP) shows the strictly conserved tyrosine Y169 and highly conserved phenylalanine F175 of human PrP (PRIO_HUMAN in figure). The sequence stretch 169-YSNQNNF-175 is required for self-assembly of the prion protein. Aromatic amino acids are shown in color. The conserved sites (*), sites with conservative (:), semi-conservative (.) and non-conservative (whitespace) replacements are annotated with symbols at the bottom. The site holding Y169 of human PrP is marked with "*" symbol and F175 with ":" symbol. Alignment of reviewed (Swiss-Prot) records of mammalian prion proteins with protein name "Major prion protein" and gene name "PRNP" or "Prnp" was performed within the Uniprot framework (Clustal-Omega with default parameters). Columns are: UniProt ID; protein name; start position; sequence; end position.

**Dataset S1.** PLAAC results per genome

**Dataset S2.** GO enrichment analysis results

**Dataset S3.** Experimentally tested cPrD list and information

**Dataset S4.** All PLAAC positive proteins and annotations

**Dataset S5.** Amino-acid codon GC and AT content

**Dataset S6.** Primers used in preparation of cPrDs for Gateway cloning

**Data and code availability:**

The code and data for reproducing the computational analyses performed for this manuscript are available at:
https://figshare.com/projects/Zajkowski_et_al_2020_Archaeal_prion_data_and_code_repository/78720
